# Co-localization of different neurotransmitter transporters on synaptic vesicles is sparse except of VGLUT1 and ZnT3

**DOI:** 10.1101/2021.06.30.449903

**Authors:** Neha Upmanyu, Jialin Jin, Henrik von der Emde, Marcelo Ganzella, Leon Bösche, Viveka Nand Malviya, Evi Zhuleku, Antonio Politi, Momchil Ninov, Ivan Silbern, Marcel Leutenegger, Henning Urlaub, Dietmar Riedel, Julia Preobraschenski, Ira Milosevic, Stefan W Hell, Reinhard Jahn, Sivakumar Sambandan

**Affiliations:** Synaptic Metal Ion Dynamics and Signaling, Max Planck Institute for Multidisciplinary Sciences, Göttingen, Germany; Department of Neurobiology, Max Planck Institute for Multidisciplinary Sciences, Göttingen, Germany; European Neurosciences Institute - A Joint Initiative of the University Medical Center Göttingen and the Max Planck Society, Göttingen, Germany; Department of NanoBiophotonics, Max Planck Institute for Multidisciplinary Sciences, Göttingen, Germany; Live-cell Imaging Facility, Max Planck Institute for Multidisciplinary Sciences, Göttingen, Germany; Bioanalytical Mass Spectrometry, Max Planck Institute for Multidisciplinary Sciences, Göttingen, Germany; Institute of Clinical Chemistry, University Medical Center Goettingen, Goettingen, Germany; Department of Structural Dynamics, Max Planck Institute for Multidisciplinary Sciences, Göttingen, Germany; Institute for Auditory Neuroscience, University Medical Center, Göttingen, Germany; Wellcome Centre for Human Genetics, Nuffield Department of Medicine, NIHR Oxford Biomedical Research Centre, University of Oxford, Oxford, UK; Multidisciplinary Institute of Ageing, MIA-Portugal, University of Coimbra, Coimbra, Portugal; Department of Optical Nanoscopy, Max Planck Institute for Medical Research, Heidelberg, Germany

**Keywords:** Synaptic vesicle, Vesicular transporters, VGLUT1, Glutamate, DyMIN STED, MINSTED, Zinc, Quantal size, Super resolution imaging, Zinc transporter 3

## Abstract

Vesicular transporters (VTs) define the type of neurotransmitter that synaptic vesicles (SVs) store and release. While certain neurons in mammalian brain release multiple transmitters, the prevalence, physiology of such pluralism and if the release occurs from same or distinct vesicle pools is not clear. Using quantitative single vesicle imaging, we show that a small population of neuronal SVs indeed contain different VTs to accomplish corelease. Surprisingly, this population is highly diverse (27 types), expressing distinct dual transporters suggesting corelease of various combinations of neurotransmitters. Using glutamatergic vesicles as an example, we demonstrate that transporter colocalization not only determine the transmitter type but also influences the SV content and synaptic quantal size. Thus, presence of diverse transporters on the same vesicle is bona-fide and, depending on the VT types, this may act as one mechanism to regulate neurotransmitter type, content and release in space and time.

## INTRODUCTION

Synaptic vesicles (SVs) are the trafficking organelles responsible for storage and release of various neurotransmitters at the synapse, with the type and content of the transmitters determined by the presence of specific vesicular transporters (VTs) in the SV membrane. Although a given neuron is thought to release only one type of neurotransmitter (referred as Dale’s principle), it is well established that certain neurons in the vertebrate central nervous system (CNS) release more than one classical (non-peptide) neurotransmitter upon activation, referred as ‘cotransmission’ (Hnasko and Edwards, 2012; Tritsch et al., 2016; Trudeau and El Mestikawy, 2018; Vaaga et al., 2014). Evidence has been mounting, especially in the last decade, that specific neurons release multiple transmitters from the same synapse (Ren et al., 2011; Shabel et al., 2014; Varga et al., 2009). While the prevalence and the physiological role of such pluralism is not clear, it has also been difficult to ascertain if the release of multiple neurotransmitters occur from the same pool of vesicles (referred as ‘corelease’) or segregated pools of SVs that may exhibit different calcium sensitivity. This knowledge is crucial for the understanding of synaptic function and computation.

Corelease of distinct neurotransmitters from the same SVs requires that the respective vesicles contain the corresponding VTs, but the available information is not sufficient to confirm the same. For both, corelease and cotransmission, the VTs must be co-expressed in the same neurons. A recent large scale single-cell transcriptomics study (involving ∼70,000 neurons) of the CNS revealed that roughly 5% of neurons co-express multiple VTs indicating the capacity for cotransmission or corelease (Zeisel et al., 2018). An analysis of the data reveals that the multi-transporter neurons are highly diverse expressing up to four different transporters, implying an unappreciated complexity of synaptic signaling (Figure S1). Considering the limitations in transcript detection in single-cell studies, it is conceivable that the actual population of neurons expressing multiple VTs may be higher, hinting widespread cotransmission or corelease in the brain.

Recent discoveries, using optogenetics, *in vivo* electrophysiology and anatomical approaches, have convincingly established the identity of an array of dual transmitter neurons and their postsynaptic partners in specific brain regions (Ren et al., 2011; Shabel et al., 2014; Takacs et al., 2018; Varga et al., 2009). The two major fast neurotransmitters, glutamate and GABA, that exert excitatory and inhibitory postsynaptic responses, respectively, have been shown to be released together with another transmitter that is of modulatory nature. For example, glutamate is released together with dopamine in the striatum and the ventral tegmental area, VTA (Chuhma et al., 2004, Stuber et al., 2010, (Hnasko et al., 2010)), or with serotonin in the hippocampal raphe nuclei projections (Sengupta et al., 2017; Varga et al., 2009), or with acetylcholine in striatal interneurons and medial habenula projections (Frahm et al., 2015; Gras et al., 2008; Higley et al., 2011; Ren et al., 2011). Similarly, GABA is cotransmitted with acetylcholine in basal forebrain projections (Saunders et al., 2015; Takacks et al., 2018), or with dopamine in the olfactory bulb (Liu et al., 2013) and the striatum (Tritsch et al., 2012). Interestingly, cotransmission of glutamate and GABA has also been observed in some neurons (Root et al., 2018; Shabel et al., 2014) including evidence for their corelease (Shabel et al., 2014). Despite this considerable progress, a compilation of recent studies on multitransmitter neurons (Table S1) reveals that only few examples exist to corroborate corelease or colocalization of transporters on the same vesicle (Fasano et al., 2017; Root et al., 2018; Saunders et al., 2015; Takacs et al., 2018; Zimmermann et al., 2015). Immunogold electron microscopy (IEM) following dual labeling of colocalized transporters provides an indication for corelease (Takacs et al., 2018; Zander et al., 2010), but it may not provide quantitative information due to limited sensitivity of IEM. Electrophysiological recordings of dual transmitter responses following optogenetic stimulations can only indicate but not conclude corelease and can only be applied to study specific pre-post neuronal populations (Shabel et al., 2014; Takacs et al., 2018). Functional *in vitro* and biochemical experiments such as measuring the effect of one transmitter on the vesicular uptake of the other (Gras et al., 2008) or SV immunosisolation (Gronborg et al., 2010) goes one notch up to explore corelease of transmitters. Nonetheless, these are bulk experiments that could lead to false-positive vesicular colocalization. Thus, the above approaches, when applied either individually or together, yield a wealth of information on dual transmitter signaling but inadequate to confirm corelease and do not address the extent of the phenomenon in the whole mammalian brain.

In the present study, we have addressed these questions by visualizing different well-characterized VTs in highly purified SVs at single vesicle level, resulting in a comprehensive and quantitative colocalization map of transporters, which reveals the extent and diversity of corelease in adult rat brain. We have also studied one example of corelease to understand the benefits and the relevance for synaptic physiology.

## RESULTS

### DyMIN STED allows high-throughput super-resolved imaging of single SVs in association with enhanced fluorescence signal

Dynamic minimum stimulated emission depletion (DyMIN STED) nanoscopy, a recently developed super resolution microscopy technique that provides super resolution with efficient fluorescence detection was used to image single SVs derived from rat brain. DyMIN STED modulates the power of the depletion light depending on the local brightness in the sample in contrast to conventional STED microscopy that applies maximum STED laser power throughout the imaging area (Gottfert et al., 2017; Heine et al., 2017). Thus, the overall light dose exposed to the sample is reduced leading to decreased photobleaching and enhanced fluorescent signal from single vesicles (Figure 1A-C). In contrast to conventional high-resolution microscopy (Farsi et al., 2016), DyMIN STED significantly improved resolution of single labeled vesicles (Figure 1B). The average size of individual SVs labeled for Synaptobrevin2 (Syb2), measured in terms of the standard deviation (SD) of an integrated Gaussian fit (σ) on each puncta in DyMIN STED was ∼33 nm in comparison to ∼126 nm in confocal, as expected for the size labeled SVs (Takamori et al., 2006). Additionally, adapting the STED laser intensity to the fluorophore density improved the signal-to-noise ratio while maintaining fluorescence signal as comparable to that of confocal microscopy (Figure 1B). To immunolabel SVs, we developed a strategy that maximized epitope coverage while effectively removing background fluorescence from unbound antibodies (Figure S2). Freshly isolated SVs of high purity (Takamori et al., 2006) were indirectly labeled using saturating concentrations of the antibody ‘in-solution’, followed by an additional size exclusion chromatography (SEC) step to remove unbound antibodies. The above method significantly increased the fluorescence brightness of single vesicles when compared to conventional labeling that is performed after immobilization of SVs on a coverslip (Figure 1D). The enhanced epitope coverage through ‘in-solution’ labeling is particularly favorable for targets with limited fluorescence budget such as VTs (Figure 1E). In addition, negative controls using samples that lacked either SVs or the primary antibody exhibited insignificant background fluorescence (∼0.1%, Figure S3A-C), which has been accounted for in all quantitative analyses. The combination of DyMIN STED and the labeling protocol thus allowed high-throughput visualization of super-resolved SVs for quantitative fluorescence analysis.

**Figure 1.**
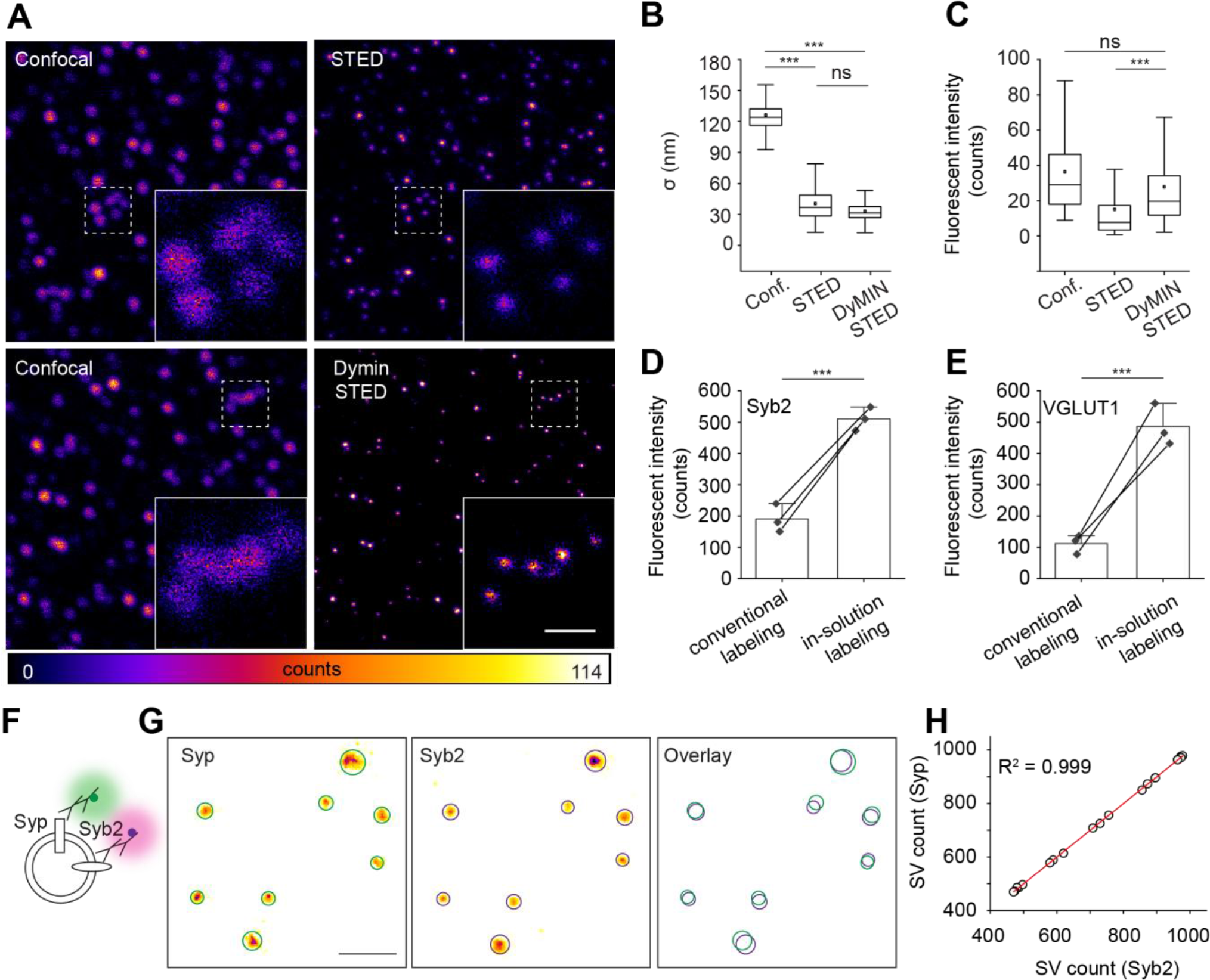
DyMIN STED allows high-throughput super-resolved imaging of ‘in-solution’ labeled single SVs in association with enhanced fluorescence signal. (**A**) Representative images of Synaptobrevin2 (Syb2) labeled SVs in confocal, conventional- and DyMIN STED mode. Insets are magnified regions marked in dashed white squares showing putative single vesicles in confocal that are clearly resolved into multiple vesicles in the conventional- and DyMIN STED. Scale bar, 500 nm. (**B**) Box plot showing a significant increase in the spatial resolution of single vesicles in the conventional- and DyMIN STED mode in comparison to confocal mode (***P<0.01; unpaired t test, n=3). (**C**) Box plot showing comparable fluorescent intensity of individual vesicles between confocal and DyMIN STED indicating enhanced fluorescence signal (reduced photobleaching) in contrast to conventional STED (p=1.74, not significant between confocal and DyMIN STED; ***P<0.01, between conventional and DyMIN STED; unpaired t-test; n=3). (**D** and **E**) Bar graphs showing significant increase in fluorescent intensity when vesicles were labeled ‘in-solution’ in comparison to conventional labeling (***P<0.01; unpaired t-test, n=3). Note that there is ∼5 fold increase in the fluorescence intensity when SVs were labeled for VGLUT1 compared to ∼2.5 fold increase for Syb2, indicating enhanced epitope coverage for low copy number proteins such as VGLUT1. (**F**) Illustration depicting double immunolabeling of purified SVs against Synaptophysin (Syp) and Syb2 using two spectrally distinct fluorescent dyes. (**G**) Representative two-color DyMIN STED images (inverse LUT) of SVs for Syp and Syb2. Circles portray SV area derived by a 2D Gaussian fit on Syp (green) and Syb2 (magenta) puncta. The overlay shows SV circle profiles of the two channels. Note the near complete overlap of all Syp and Syb2 circles. Scale bar, 500 nm. (**H**) Scatter plot of the detected Syp and Syb2 puncta count in different experiments. Linear regression of the data shows even distribution of Syp and Syb2 (n = 4 experiments, >14,000 SVs for both channels). See also Figures S2, S3 and S4.

### Two-color DyMIN STED reveals colocalization pattern of diverse transporters on the same vesicle

Two-color DyMIN STED imaging of Synaptophysin (Syp), a marker present on all SVs irrespective of the neurotransmitter phenotype ((Takamori et al., 2000), Figure S4) and Synaptobrevin2 (Syb2) showed colocalization of nearly all puncta in both channels revealing that both Syp and Syb2 act as a marker for total number SVs (Figure 1F-H). Colocalization was determined using a co-ordinate based colocalization analysis ((Malkusch et al., 2012), Figure S4 and S5, STAR Methods) since SV puncta followed a Rayleigh distribution as described in previous reports (Geumann et al., 2008). To determine the proportion of SVs expressing a given transporter within the total SV population, we performed dual immunolabeling of SVs for Synaptophysin (Syp) and each of the following vesicular transporters - VGLUT1, VGLUT2, VGLUT3 (specific for glutamate), VGAT (specific for GABA and Glycine), VMAT2 (specific for monoamine transmitters), VAChT (specific for acetylcholine), and ZnT3 (specific for Zn^2+^, Figure 2A). VGLUT1, VGLUT2 and VGLUT3 were detected on approximately 60%, 23% and 2.8% of all SVs respectively, in agreement with previous reports ((Takamori et al., 2006; Wilhelm et al., 2014), Figure 2A). VGAT labeling was detected on 15.9% of all SVs, again confirming earlier reports (Takamori et al., 2006; Takamori et al., 2000). VMAT2 and VAChT were present on 2.6% and 1.5% of all SVs, respectively (Figure 2A). Surprisingly, ZnT3, the transporter responsible for loading Zn^2+^ that inhibits post synaptic AMPA and NMDA receptors (Kalappa et al., 2015; Palmiter et al., 1996; Romero-Hernandez et al., 2016) (Palmiter et al., 1996; Romero-Hernandez et al., 2016), is one of the most abundant of all VTs; a staggering ∼30% of all SVs were ZnT3 positive (Figure 2A). Indeed, ZnT3 protein has been consistently detected in earlier SV proteomics studies (Morceano et al., 2005, Takamori et al., 2006, Grönborg et al., 2010, Boyken et al., 2013) and also been shown to preferentially co-express at excitatory synapses when compared to inhibitory ones (Grönborg et al., 2010, Palmiter et al., 1996). Although the VT antibodies used above are well characterized (Gras et al., 2008; Herzog et al., 2006; Hnasko et al., 2010; Takamori et al., 2000), we tested their specificity using purified chromaffin granules (CG, Figure S3D), which share many proteins with SVs including the fusion machinery but lack the above VTs. Labeling and single CG imaging was performed as described above for SVs. None of the VT antibodies resulted in staining, confirming their high specificity (Figure S3). The marginal background fluorescence primarily due to secondary antibodies (∼0.1%) was accounted for in all single vesicle quantifications.

**Figure 2.**
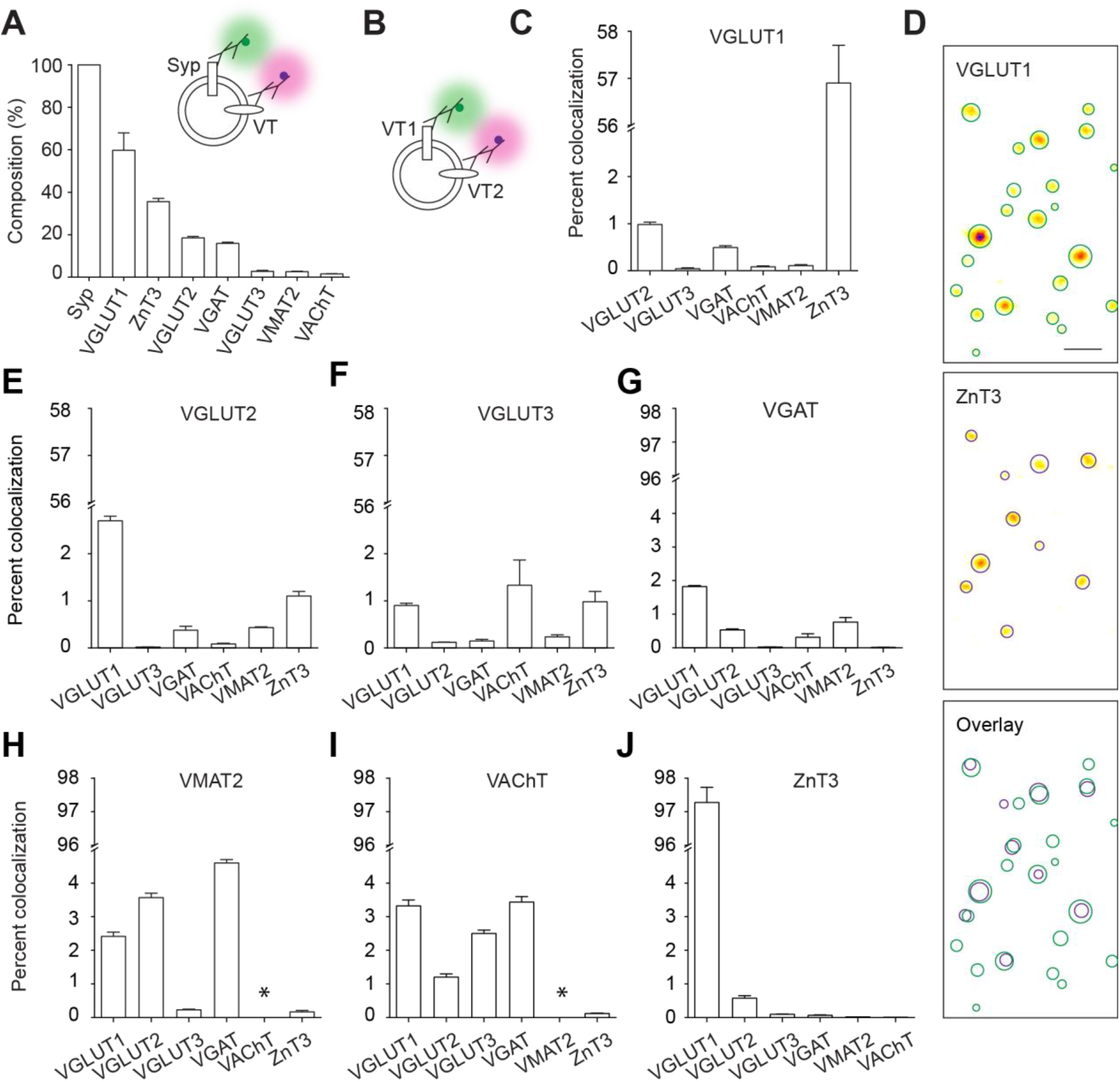
Two color DyMIN STED reveals colocalization pattern of diverse transporters on the same vesicle. (**A**) Bar graph quantifying the proportion of SVs expressing a specific VT. Values are normalized to Syp, which acted as reference for total SVs. The inset illustrates double immunolabeling of SVs against Syp and a VT (n = 3 experiments for each pair; >10000 total SVs for each dual labeling). (**B**) Illustration depicting dual immunolabeling of SVs against two different VTs (VT1 and VT2). (**C)** Bar graph showing the degree of vesicular colocalization of VGLUT1 against the specific VTs on the X-axis (n > 3 experiments for each pair; >10000 total SVs for each dual labeling). (**D**) Representative images (inverse LUT) of VGLUT1 and ZnT3 channels from the two-color DyMIN STED experiment shown in C. Circles portray the SV area derived by a 2D Gaussian fit on VGLUT1 (green) and ZnT3 (magenta) puncta. The overlay shows SV circle profiles of the two channels (n > 3 experiments; >20000 total SVs for both VGLUT1 and ZnT3). Scale bar, 200 nm. (**E-J**) Bar graphs showing the degree of vesicular colocalization of a specific VT indicated on top of the each graph against the VTs on the X-axis (n = 3 experiments for each pair; >10000 total SVs for each dual labeling). No data are shown for VMAT2 and VAChT combination. All values are means ± SEM. See also Figure S3. S4, S5 and S6.

To investigate expression of multiple VTs on the same vesicle, we measured the degree of colocalization between two distinct transporters in all possible pairwise combinations using dual color DyMIN STED microscopy (Figure 2B-J). For glutamatergic vesicles (excitatory), interesting differences became apparent when comparing the three VGLUT isoforms (VGLUT-1-3). First, the overlap between the two abundant variants VGLUT1 and VGLUT2 was low (Fig 2H and I, ∼1% of VGLUT1 SVs contain VGLUT2) in contrast to their coexistence during development (Herzog et al., 2006) but in agreement with the non-overlapping expression pattern of the two VTs in adulthood (El Mestikawy et al., 2011; Takamori et al., 2001). Second, while VGLUT2 did not show any preference to dopaminergic or cholinergic vesicles, VGLUT3 showed preference to VAChT (Figure 2E and F). On the other hand, VMAT2 showed preference to VGLUT2 whereas VAChT showed no such preference to any VGLUTs (Figure 2H and I). Coexpression of the above VTs was previously observed in dopaminergic and cholinergic neurons in the midbrain and the striatum (Gras et al., 2008; Hnasko et al., 2010). Third, VGLUT1 but not the two other variants showed preferred colocalization with ZnT3 (∼57%, Fig 2C-E and J). We confirmed the robust colocalization of ZnT3 and VGLUT1 in SV preparations from mouse using three different antibodies of ZnT3, for which, specificity was validated by western blot or single vesicle imaging using using ZnT3 knockout (KO) mice (Figure S6).

VGAT, the transporter defining GABAergic vesicles, was shown previously to be colocalized with other VTs at many synapses (Tritsch et al., 2016). However, on the vesicle level, VGAT shows marginal overlap with other VTs. For instance, VMAT2 and VAChT are only found on ∼0.75% and ∼0.3% of GABAergic vesicles, respectively (Figure 2G), suggesting that corelease of GABA with monoamines or acetylcholine from the same vesicle pool is not widespread. On the contrary, both monoaminergic and cholinergic SVs generally showed marginal degrees of colocalization with VGLUTs and VGAT (Figure 2H and I, note that colocalization between VMAT2 and VAChT could not be measured, STAR Methods). Finally, we also examined SVs that carry one of the VGLUT variants and VGAT. Both VGLUT1 and VGLUT2 showed similar levels of colocalization with VGAT (slightly below 0.5%). Overall, VTs exhibit a diverse combination of colocalization on single SVs.

### SVs are highly diverse based on the phenotype of expressed vesicular transporters

The above colocalization pattern of VTs and the proportion of SVs carrying a specific VT in the entire SV population allowed us to quantify the relative abundance of SVs carrying specific VT phenotype. (Figure 3A and B). We identified 27 classes of SVs, each carrying either one or two distinct VTs (Figure 3A). Interestingly, these SVs classes sum up to ∼102% indicating coverage of the total SV population by the labeling method. This is also consistent with a recent finding that SVs contain only previously recognized *bona fide* neurotransmitters (Chantranupong et al., 2020). The fraction of SVs specific for a single neurotransmitter (i.e. carrying only one VT) is ∼64% suggesting that roughly one-third of SVs are capable of storing more than one neurotransmitter. The multitransmitter SV population, however, is predominated by VGLUT1-ZnT3 vesicle, which accounts for ∼34% of all SVs. To validate such a high degree of colocalization between VGLUT1 and ZnT3, we performed IEM of isolated SVs using VGLUT1 and ZnT3 primary antibodies and secondary species conjugated to gold nano particles (Figure 3A inset, VGLUT1-5 nm, ZnT3-10 nm). Indeed, a modest population of SVs was positive for both VGLUT1 and ZnT3, in spite of the lower sensitivity of IEM when compared to light microscopy.

**Figure 3.**
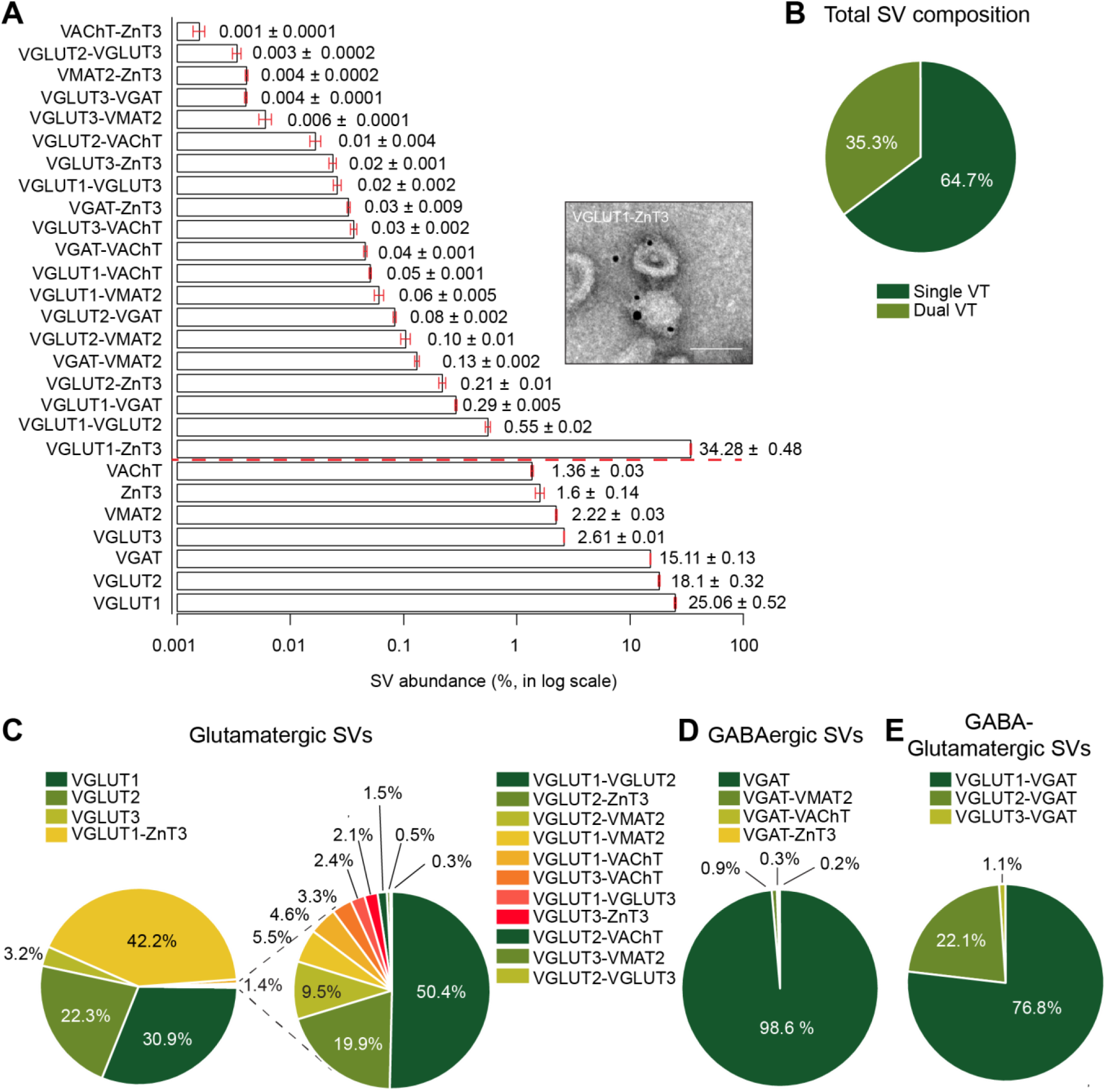
SVs are highly diverse based on the phenotype of expressed vesicular transporters. (**A**) Bar graph showing the relative abundance of different vesicle classes containing either one (below dashed red line) or two distinct VTs (above the red dashed line), obtained from colocalization degree of each VT pairs and the respective proportion of each VT in the total SV pool as shown in Figure 2. The numbers next to each bar denotes the actual value. Inset-representative electron immunogold microscopy (IEM) image validating colocalization of VGLUT1 and ZnT3 by displaying gold nano-particles specific for VGLUT1 (5 nm) and ZnT3 (10 nm) (n = 2 IEM experiments, 4.14% of all SVs carried gold particles specific for both VGLUT1 and ZnT3; 35.6% and 6.7% of vesicles were positive for VGLUT1 and ZnT3 alone; negative controls that lacked primary antibodies exhibited negligible signals). Bar graph values are means ± SEM. Scale bar in the inset, 50 nm (**B**) Pie diagram showing broad composition of total SVs containing either one or two distinct VTs. (**C**) Left, pie diagram showing the composition of glutamatergic vesicles expressing either one of the three vesicular glutamatergic transporters (VGLUT1-3) or in association with another VT. Right, the small components are visualized by an exploding pie chart recalibrated to 100% for greater visibility. (**D**) Similar pie chart, as shown in C, for GABAergic vesicles expressing VGAT. (**E**) Similar pie chart, as shown in C, for SVs expressing one of the three VGLUTs and VGAT. All statistics above were obtained from two color DyMIN STED images. See also Figure S1

We also determined the composition of excitatory SVs, characterized by the expression of either one of the three VGLUT subtypes (VGLUTs) or in combination with another VT (Figure 3C). While a variety of excitatory SVs involving different combinations of VGLUTs and other transporters exist, two defining features are very conspicuous – 1) the sheer abundance of VGLUT1-ZnT3 SVs (∼42% of excitatory SVs), which we will focus in the later sections and that 2) only ∼0.7% of the excitatory SVs carry both VGLUT1 and VGLUT2. The low overlap between VGLUT1 and VGLUT2, in spite of their co-expression during early development stages (Herzog et al., 2006), reflects a near complete exchange of VGLUT1 for VGLUT2 in the adult cortex.

Inhibitory SV composition, characterized by the expression of only VGAT or in combination with another VT, has also been deduced (Figure 3D). Interestingly, only 3.5% of VGAT positive SVs carry another transporter, which is not anticipated in view of the increasing evidence for cotransmission of GABA together with other neurotransmitters (Tritsch et al., 2016) suggesting that VGAT and other VTs exist in segregated pools at those synapses. A very small population of GABA-Glutamatergic SVs (∼0.37% of all SVs) has also been detected carrying VGAT and one of the VGLUT subtypes. Interestingly, the composition of GABA-Glutamatergic SVs also provides some insight on the VGLUT subtype that is majorly involved in GABA corelease (Figure 3E). Among the three subtypes, VGLUT1 is preferentially colocalized with VGAT, although only VGLUT2 has been predominantly detected for corelease of glutamate with GABA (Shabel et al., 2014).

Taken together, the synaptic vesicles carrying two VTs are highly diverse although the overall degree of colocalization between any transporter pair is low (consistently below 5% of all SVs), which is in consistent with the abundance of neurons capable of releasing more than one neurotransmitter (Figure S1). Thus, a vast majority of all SVs is indeed specific for only a single neurotransmitter, with an exception of VGLUT1-ZnT3 SVs.

### ZnT3 is a highly abundant SV protein specifically enriched in VGLUT1 positive vesicles

ZnT3 has been detected as an SV protein in earlier studies, especially enriched in VGLUT1 positive synapses. However, the relative abundance of ZnT3 positive vesicles and its colocalization degree with VGLUT1 is unexpected. To verify the high abundance of VGLUT1-ZnT3 SVs, we conducted a series of quantitative investigations to deduce ZnT3 protein abundance, expression and distribution. First, we measured the amount of ZnT3 protein in SVs using quantitative immunoblotting using highly purified recombinant ZnT3 protein (42 kDa) as standard (Figure 4A and B). Increasing amounts of SVs and purified ZnT3 were loaded on the same gel and ZnT3 was detected using a specific monoclonal antibody (Figure 4A). The amount ZnT3 in the SVs, obtained from the standard curve, was ∼12 ng/µg of SVs (Figure 4B, 262±23 fmol/µg of SVs), revealing that ZnT3 is one of the most abundant SV membrane proteins (∼1.5% of the total amount of SV proteins, see (Takamori et al., 2006)). To validate the above quantification, we performed quantitative mass spectrometry (MS) of the SV proteome using a label-free intensity-based absolute quantification (iBAQ) approach (Figure 4C). We detected ∼1500 proteins including low abundant transporters such as VMAT2 and VAChT, each with individual IBAQ values (Table S2). In agreement with immunoblotting measurements, ZnT3 was detected as an abundant SV protein with approximately one-third of absolute amount of VGLUT1 (Figure 4C). To understand how this differential protein amount of ZnT3 and VGLUT1 reflects as copy number per vesicle, we estimated the copy number of the proteins by dividing the number of molecules by the number of ZnT3 positive vesicles per µg of SVs (arrived from (Takamori et al., 2006) and Figure 2D). The average ZnT3 copy number was estimated to be roughly half of the average copy number of VGLUT1 (Figure 4C).

**Figure 4.**
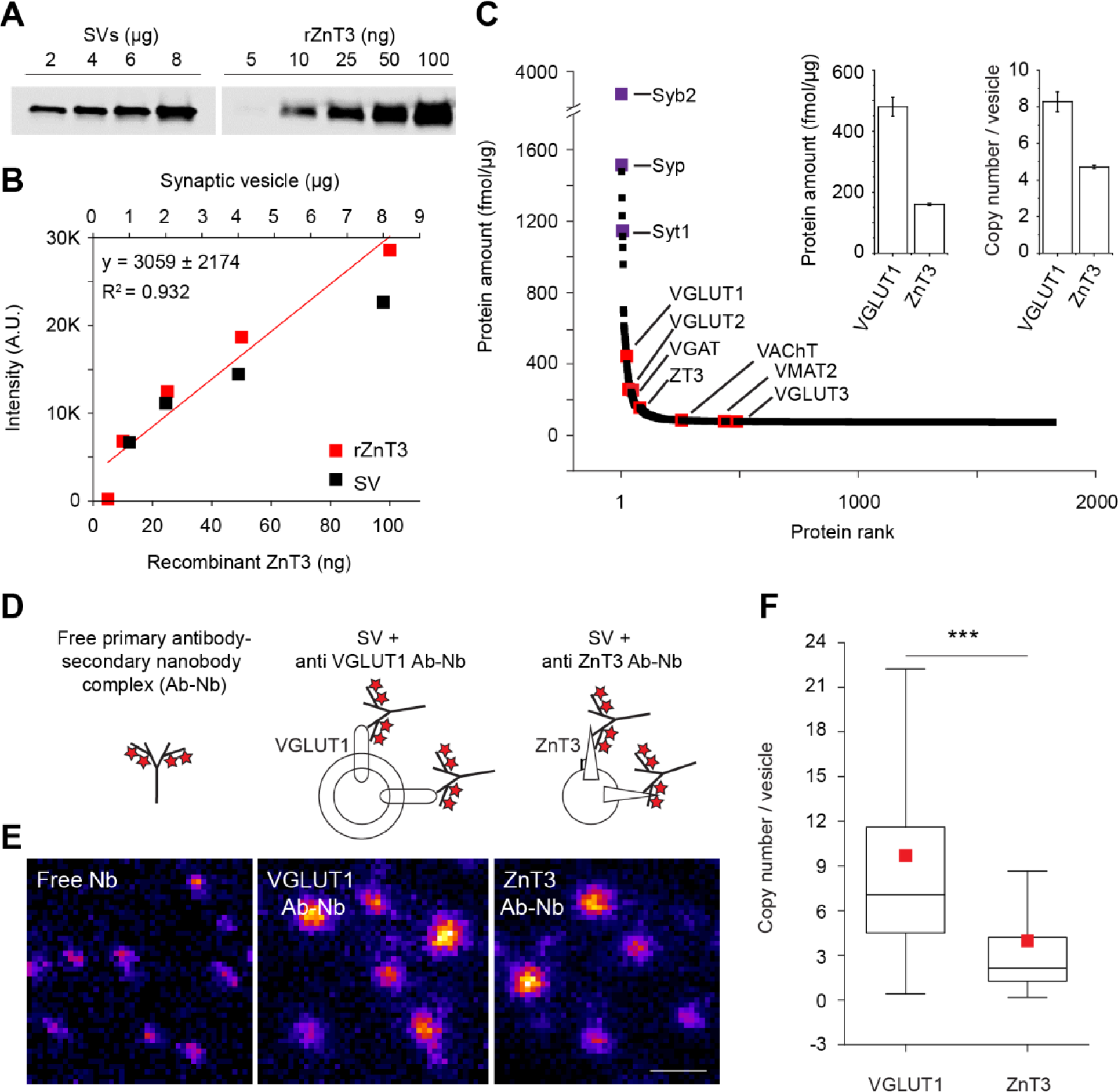
ZnT3 is a highly abundant SV protein specifically enriched in VGLUT1 positive vesicles. (**A**) An example quantitative immunoblot for ZnT3 in SVs using purified recombinant ZnT3 protein as standard (rZnT3). Highly purified SVs and rZnT3 were loaded onto the same gel at specified concentrations. (**B**) Scatter plot for quantification of ZnT3 abundance in SVs following quantitative immunoblotting using the standard curve for rZnT3 fitted with a linear function (n=2 biological samples). (**C**) SV proteins ranked by their abundance as estimated using quantitative mass spectrometry (MS). Selected SV proteins are color-coded for visibility; purple for specified SV markers (Syb2-synaptobrevin2; Syp-synaptophysin; Syt1-synaptotagmin1) and red for the VTs investigated in the current study (VGLUT1, VGLUT2, ZnT3, VGLUT3, VGAT, VMAT2 and VAChT). Inset: Bar graphs showing relative abundance (left) and copy number (right) of VGLUT1 and ZnT3 estimated from MS analysis (n=2 biological samples). (**D**) Illustration of purified SV labeling to preserve the stoichiometric ratio between VGLUT1 (middle) or ZnT3 (right) and the number of fluorophores using monoclonal primary antibody and secondary nanobody conjugated to fixed number of fluorophores (Ab-Nb). The left illustration shows free Ab-Nb complex. (**E**) Representative DyMIN STED images (fire) of free Ab-Nb (left) and SVs labeled against VGLUT1 (middle) and ZnT3 (right) as described above. Scale bar, 100 nm. (**F**) Box plot quantifying the copy numbers of VGLUT1 and ZnT3 in individual SVs (n=3, p<0.001, unpaired t-test).

To directly measure the copy number of VGLUT1 and ZnT3 on individual SVs, we developed an imaging based approach using DyMIN STED (Figure 4D-F). Isolated SVs were labeled either for VGLUT1 or for ZnT3 using primary monoclonal antibody and secondary nanobody (Sograte-Idrissi et al., 2020) conjugated to a definite number of fluorophores resulting in a fixed stoichiometry between the number of protein molecules and fluorophores. Following single vesicle imaging, the copy number was obtained by dividing the SV fluorescence signal with the mean signal of the free antibody-nanobody complex (Figure 4E and F, STAR Methods). Compared to conventional estimates of average copy numbers that rely on bulk measurements (Takamori et al., 2006; Taoufiq et al., 2020), single vesicle imaging based approaches reveal the variability between vesicles (Mutch et al., 2011). While the VGLUT1 and ZnT3 copy numbers were highly variable between the vesicles, the average copy number of ZnT3 was significantly lower than that of VGLUT1 (VGLUT1, 9.46±.06 (mean±SEM), 7.96 (SD); ZnT3, 3.63±.07 (mean±SEM), 2.45 (SD), Figure 4F), but consistent with the above MS estimations.

Finally, to corroborate colocalization of ZnT3 with VGLUT1, we performed immunoisolation of ZnT3 SVs using a monoclonal anti-ZnT3 antibody and detected selected proteins through western blotting. ZnT3 immunoisolated SVs showed, as expected, selective enrichment of VGLUT1 when compared to the control SVs that are immunoisolated using synaptobrevin2 antibodies representing all SVs (Figure S7). In addition to VGLUT1, another SV membrane protein, SV2B was also enriched in ZnT3 sample concurring with earlier findings in SVs immunoisolated with VGLUT1 antibodies (Gronborg et al., 2010). Overall, the above data reveals that ZnT3 is an abundant SV protein that is broadly expressed as an integral component of VGLUT1-containing SVs.

### Heterogeneous pools of SVs containing VGLUT1, ZnT3 and VGLUT1-ZnT3 reside at a single synapse

VGLUT1 and ZnT3 exhibit an overlapping expression pattern on the mesoscale (McAllister and Dyck, 2017). To compare the synaptic distribution of VGLUT1 and ZnT3 SVs, we labeled the transporters together with an active zone marker, Piccolo, in cultured primary hippocampal neurons and visualized them using confocal microscopy. A pixel based correlation of diffraction limited images revealed a high degree of overlap between both transporters (Figure S6E and G) indicating that almost every VGLUT1 synapse is positive for ZnT3. To further probe the intrasynaptic distribution of VGLUT1 and ZnT3, we imaged individual synapses at single fluorophore level using MINSTED nanoscopy (Figure 5A-M). MINSTED is the most recent addition to the STED superresolution family, carrying out the DyMIN STED principle on individual fluorophores with a highly efficient sampling algorithm and thus ultimately providing spatial resolution at the molecular scale (1-3 nm) (Weber et al., 2021). To achieve stochastic on/off switching of emitters, we performed DNA PAINT (Schnitzbauer et al., 2017) by labeling VGLUT1 and ZnT3, each with a specific ssDNA oligo (docking strand), through indirect immunolabeling. The transient binding of a complementary strand coupled to a fluorescent label (imager strand) in the imaging buffer was harnessed to capture stochastic fluorescence emission. The Piccolo signal marked the active zone (AZ) and was acquired in confocal mode in a large field of view followed by two-color MINSTED imaging in selected synaptic regions. Multiplexing was performed by sequential image acquisition with two different imager strands containing distinct oligo sequences (Schnitzbauer et al., 2017). Overall, the MINSTED-PAINT combination kept the background fluorescence low while achieving a localization precision of fluorophores below 1.5 nm in the focal plane. Individual vesicles, identified by cluster analysis of the binding events, were characterized for size, nearest neighbor distance, number of vesicles per AZ, distance to the AZ and the number of binding events per vesicle (Figure 5A-H). The characterization of VGLUT1 positive vesicles correlates with the ultrastructural organization of vesicle clusters obtained through high pressure fast freezing electron microscopy (Borges-Merjane et al., 2020).

**Figure 5.**
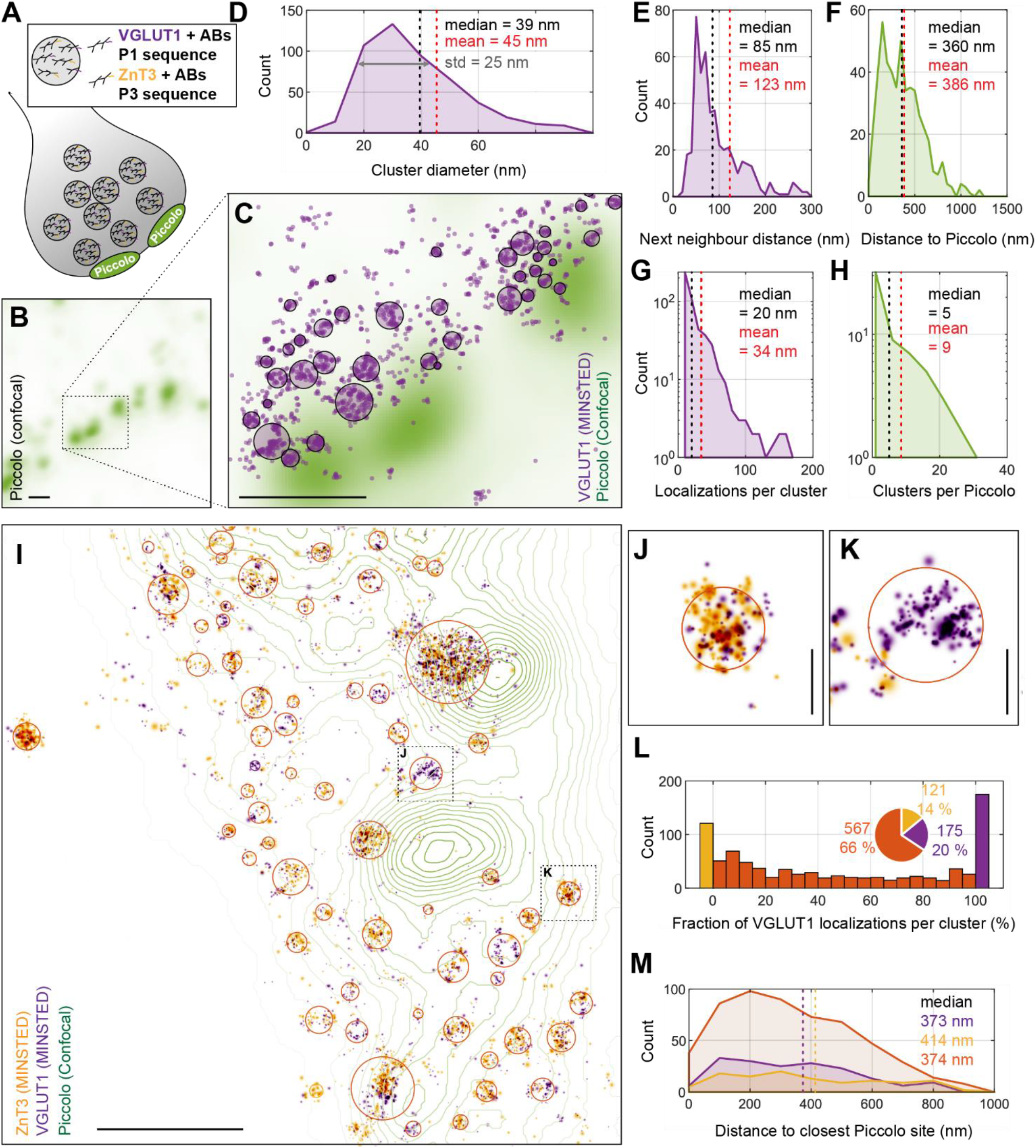
Heterogeneous pools of SVs containing VGLUT1, ZnT3 and VGLUT1-ZnT3 reside at individual synapses. (**A**) Illustration showing DNA PAINT labeling of VGLUT1 and ZnT3 with Piccolo as AZ marker in hippocampal culture neurons. (**B**) A representative confocal image of a selected synapse region showing Piccolo distribution. Scale bar, 400 nm. (**C**) An image of the zoomed-in region from B showing MINSTED localization position estimates of VGLUT1 with Piccolo in green background. The violet circles indicate the two-sigma range around the VGLUT1 cluster centers. Scale bar, 400 nm. (**D)** Size distribution of VGLUT1 clusters with a bin size of 10 nm. The cluster diameter is estimated by 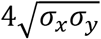, where *σ*_*x*_ and *σ*_*y*_ denote the standard deviation of the assigned localizations in x- and y-direction. (**E)** Distance distribution VGLUT1 clusters to its nearest neighbor with a binsize of 10 nm. (**F)** Distance distribution of VGLUT1 clusters to its nearest AZ with a bin size of 50 nm. AZ centers were defined from the Piccolo confocal images as local maxima with a minimum prominence of 30% with respect to the global maximum. **(G)** Distribution of the number of assigned localizations per VGLUT1 cluster with a bin size of 10. (**H)** Distribution of the number of VGLUT1 clusters assigned to each Piccolo site with a bin size of 5 (n=4 experiments, 59 AZ and 504 VGLUT1 clusters for D-H) (**I)** A representative image showing a rendered two-color MINSTED image of VGLUT1 and ZnT3 localizations with a pixel size of 0.3 nm. Each localization is displayed as a Gaussian with a standard deviation given by its estimated localization precision. For better visibility of highly precise localizations, an offset is set at 0.5 nm. The respective confocal Piccolo channel is shown as an overlaid contour plot. The two-sigma range around each cluster’s center position is indicated by a circle. Scale bar, 400 nm. (**J and K)** Representative VGLUT1 clusters marked in I rendered at full precision showing contrasting ZnT3 localizations. Scale bar, 50 nm. (**L)** Distribution of the fraction of VGLUT1 localizations per cluster at a bin size of 5 %. Clusters with 100% VGLUT1 localizations are defined ‘VGLUT1 only’ vesicle (Magenta); Clusters with 100% ZnT3 localizations are defined as ‘ZnT3 only’ vesicles; all intermediate fractions are assigned as ‘colocalized SVs’ containing both VGLUT1 and ZnT3. (**M)** Distribution of the distances of the above three SV types to the next Piccolo site with a bin size of 100 nm (n=4 experiments, 59 AZ and 863 clusters for L-M). See also Figure S6.

MINSTED DNA PAINT multiplexing directly revealed that ZnT3 vesicles colocalize with VGLUT1 containing SVs in individual synapses (Fig 5I-L). We visualized three different pools of SVs carrying distinct VT types at individual synapses - only VGLUT1, only ZnT3 and VGLUT1-ZnT3. While, on average, there is no significant difference in the number of vesicles positive for VGLUT1 or ZnT3 per synapse, vesicles carrying both VGLUT1 and ZnT3 accounted for roughly 66% of all SVs, which is significantly higher than the whole brain statistics obtained from purified SVs (Figure 3A), suggesting enrichment of VGLUT1-ZnT3 SVs in the hippocampus (Fig 5L). We found no difference between the three different SV pools with their respective lateral distances to AZ (Fig 5M). The current study reveals VGLUT1-ZnT3 SVs as the single largest pool at the synapse, presumably due to similar trafficking pathways for VGLUT1 and ZnT3 (Salazar et al., 2005).

### Zn^2+^ facilitates vesicular glutamate uptake by a mechanism mediated by ZnT3

Does the presence of an active Zn^2+^-transport affect glutamate transport activity? To address this issue, we measured glutamate uptake in enriched SV fractions (LP2, (Preobraschenski et al., 2014)) using a purified protein-based glutamate sensor, iGluSnFR (Marvin et al., 2018) in the presence of different Zn^2+^ concentrations, ([Zn^2+,^], Figure 6A-C). Glutamate uptake was performed using 4 mM ATP, 4 mM KCl, 10 mM potassium glutamate in MOPS-Glycine uptake buffer in the presence or absence of proton ionophore, FCCP, and high affinity specific Zn^2+^ chelator, TPEN. The data were corrected for uptake in the presence of FCCP and normalized to the TPEN condition. Glutamate uptake into vesicles was dose-dependently facilitated by Zn^2+^ and exhibited a ∼2 fold increase in the maximal uptake with an EC50 of ∼25 nM (Figure 6B and C), a physiologically relevant cytoplasmic [Zn^2+^] in view of the confined volume of a typical presynaptic compartment (3.7 x 10^-22^ L, (Goch and Bal, 2020; Wilhelm et al., 2014)) and activity-dependent increase of cytoplasmic [Zn^2+^] (Sanford et al., 2019). To determine if the facilitatory Zn^2+^ effect is mediated by ZnT3, we performed the glutamate uptake assay in preparations from ZnT3 KO mice. Although KO mice showed no apparent change in VGLUT1 expression level (Figure S6), the basal glutamate uptake performed in the presence of 1 pM Zn^2+^ was slightly reduced (∼20%, Figure 6A) suggesting that ZnT3 is involved in the determination of glutamate quantal size in resting conditions concurring with the reduction in the amplitude of miniature and spontaneous excitatory postsynaptic currents observed in ZnT3 KO mice (Lavoie et al., 2011). Moreover, the facilitatory effect of Zn^2+^ on glutamate uptake was completely abolished in KO implying the existence of an activity-dependent regulation of vesicular glutamate content by Zn^2+^ through ZnT3 (Figure 6B).

**Figure 6.**
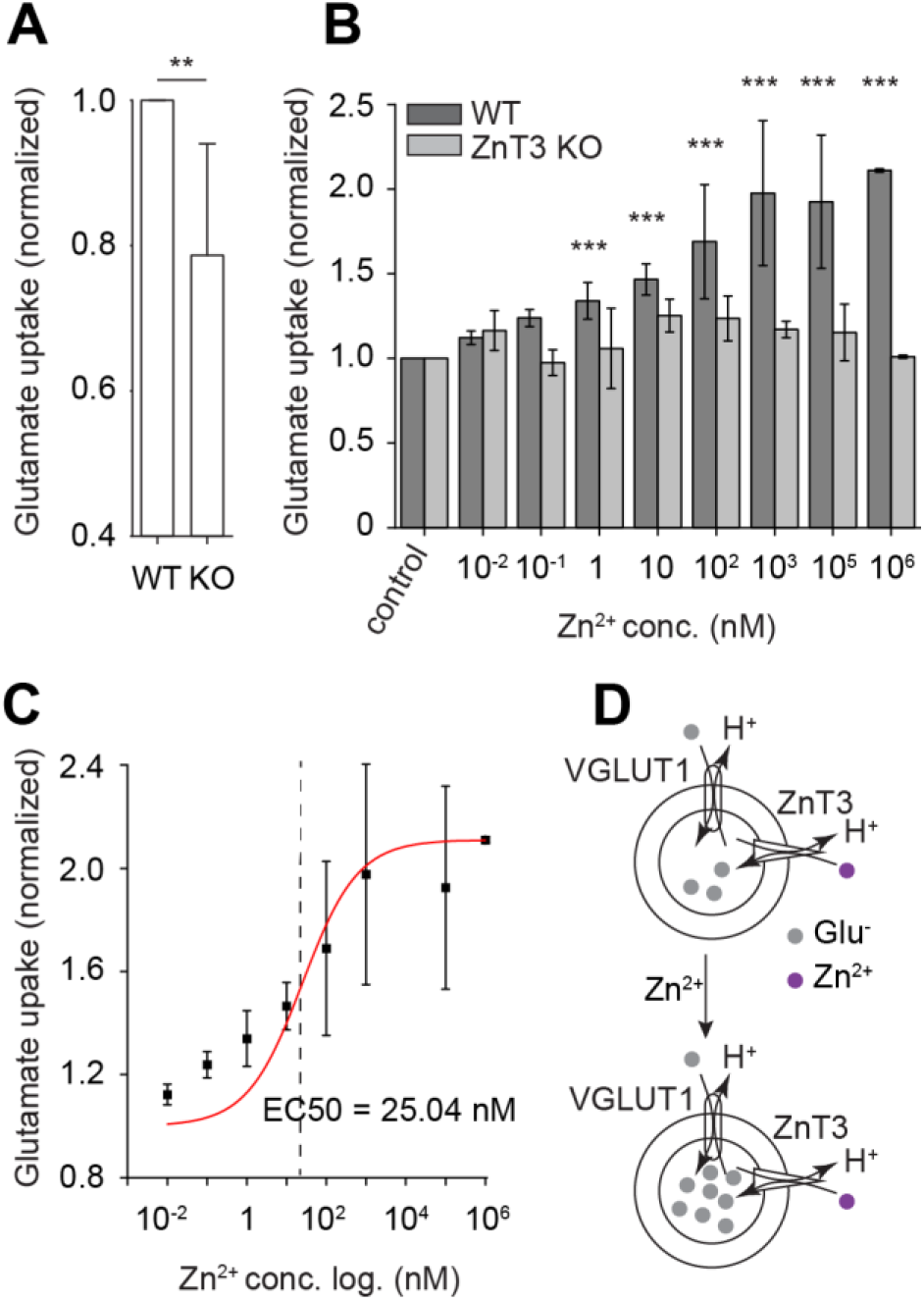
Zn^2+^ facilitates vesicular glutamate uptake by a mechanism mediated by ZnT3. (**A**) Bar graph showing enhanced basal vesicular glutamate uptake in isolated SVs (LP2 fraction) in ZnT3 wildtype mice (WT) in comparison to knockout (KO, *P<0.05, n=6 experiments, paired *t*-test). (**B**) Bar graph showing enhancement of glutamate uptake in increasing concentrations of Zn^2+^ in WT, which is absent in SVs prepared from ZnT3 KO mice (***P<0.001, n=6 experiments, two-way ANOVA). (**C**) Scatter plot showing dose-dependent facilitation of glutamate uptake by Zn^2+^. The data points were fitted with a sigmoidal function (Logistic5) to calculate the EC50 value. (**D**) Illustration of enhanced vesicular glutamate uptake by Zn^2+^. All values are means ± SEM.

### Ambient Zn^2+^ increases synaptic quantal size via ZnT3

To determine if the facilitatory effect of Zn^2+^ on vesicular glutamate uptake lead to enhanced quantal glutamate release, we explored if Zn^2+^ affects miniature excitatory postsynaptic currents (mEPSCs), which represents synaptic quantal release of glutamate. We performed whole-cell recordings in hippocampal CA3 pyramidal neurons, which receives signals from ZnT3 enriched mossy fiber terminals. AMPAR mediated mEPSCs were obtained in the presence of TTX, bicuculline and D-APV in ZnT3 wildtype (WT) and KO mice. As reported previously (Lavoie et al., 2011), the mEPSC peak amplitude and frequency were decreased in ZnT3 KO animals when compared to the WT (Figure 7B-D). Interestingly, the level of reduction in the peak amplitude (∼20%) is comparable to the magnitude of reduction in glutamate uptake in ZnT3 KO SVs (see Figure 6A) suggesting that ZnT3 mediated facilitation of vesicular glutamate uptake underlies the basal synaptic quantal size in the WT. The reduction in the frequency is also consistent with earlier findings of higher calcium sensitivity of SVs in the WT (Lavoie et al., 2011). Next, to determine if Zn^2+^ mediated facilitation of vesicular glutamate uptake leads to enhanced quantal size, we explored the effect of exogenous application of Zn^2+^, which results in increased cytoplasmic Zn^2+^ concentration (Sanford et al., 2019), on quantal properties. We measured mEPSCs in the presence of 20 µM of Zn^2+^ in the bath solution following a baseline recording. There was no change in mEPSCs following Zn^2+^ application (n=8, data not shown). Since Zn^2+^ mediated facilitation of glutamate uptake is dose dependent, we performed the above experiment in higher Zn^2+^ concentration. In the presence of 200 µM of Zn^2+^, the mEPSC peak amplitude was significantly increased in the WT (Figure 7B-D) indicating Zn^2+^ mediated regulation of glutamatergic quantal size. However, there was no change in the frequency, decay and rise time kinetics indicating that ambient Zn^2+^ did not alter release probability or induce postsynaptic changes. Finally, we tested the effect of ambient Zn^2+^ on neurotransmission in ZnT3 KO mice. No changes in the e in the mEPSC amplitude or frequency (Figure 7B-D) was detected in the KO indicating that the increase in glutamate quantal size by the ambient zinc is mediated via ZnT3 and not due to endocytic uptake of zinc. Overall, the above data demonstrate dose-dependent regulation of glutamatergic neurotransmission by enhancing synaptic quantal size via ZnT3 co-present on VGLUT1 SVs.

**Figure 7.**
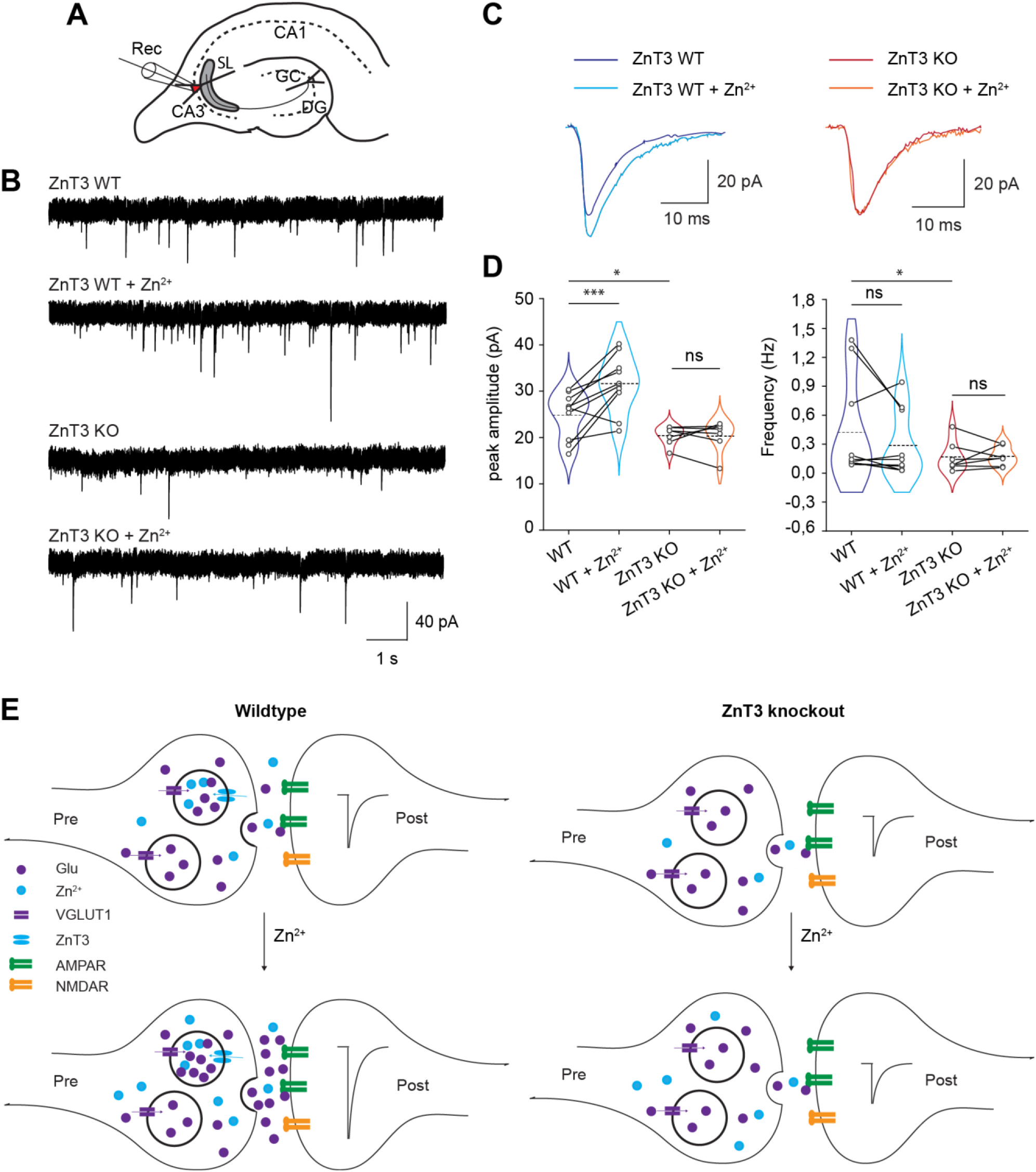
Ambient Zn^2+^ increases synaptic quantal size via ZnT3. (**A**) Scheme of whole-cell recordings performed on hippocampal CA3 neurons. (**B**) Representative traces of mEPSC activity during baseline recording and after Zn^2+^ application in ZnT3 WT and KO mice. (**C**) Average traces showing time-course of mEPSCs during baseline and after Zn^2+^ application in WT (left, decay, 6.6±0.4 ms during baseline vs 7.48±0.7 ms after Zn^2+^, p>0.2, rise time, 1.0±0.16 ms during baseline vs 1.1±01 ms after Zn^2+^, p>0.5, unpaired t-test) and KO (right, decay: 7.8±1.2 ms during baseline vs 6.8±0.7 ms after Zn^2+^, p>0.20 unpaired t-test; rise time: 1.15±0.14 ms during baseline vs 1.1±0.1 ms after Zn^2+^, p>0.5, unpaired t-test). (**D**) Violin plots comparing the peak amplitude (left) and frequency (right) of mEPSCs between baseline recordings and after Zn^2+^ application in ZnT3 WT and KO mice. Left: The baseline peak amplitude of mEPSCs was decreased in the KO compared to WT mice (24.9±1.5 pA in WT, n=10; 20.73±0.7 pA in KO, n=7; *p<0.05, t-test). Following Zn^2+^ application, the peak amplitude significantly increased in the WT (24.9±1.5 pA during baseline; 31.2±1.9 pA after Zn^2+^ application, n=10, ***p<0.002, unpaired t-test) and not in the KO mice (20.7±0.7 pA during baseline; 20.4±1.2 pA after Zn^2+^ application, n=7, p>0.5, unpaired t-test). Right: The baseline frequency was slightly reduced in the WT when compared to KO (WT, 0.42±0.16 Hz, n=10; KO, 0.16±0.06 Hz, n=7, p<0.05, unpaired t-test). Following Zn^2+^ application, the frequency did not change in both WT (0.42±0.1 Hz during baseline; 0.28±0.1 after Zn^2+^, p>0.1, unpaired t-test, n=10) and KO (0.16±0.06 during baseline; 0.17±0.03 after Zn^2+^ application, n=7, p>0.5, unpaired t-test). (**E**) Illustration showing the significance of colocalization of VGLUT1 and ZnT3 on the same synaptic vesicles in the regulation of glutamate quantal size during synaptic activity. Top left: colocalization of VGLUT1 and ZnT3 leads to co-uptake and corelease of glutamate and Zn^2+^ from same SV pools in resting conditions. Bottom left: In the presence of ambient Zn^2+^, more glutamate is taken up into SVs expressing both VGLUT1 and ZnT3 leading to enhanced glutamate release and postsynaptic response. Top right: genetic deletion of ZnT3 leads to reduced vesicular glutamate content and postsynaptic response at resting conditions. Bottom right: Absence of ZnT3 does not affect vesicular glutamate content or postsynaptic response in the presence of ambient Zn^2+^ demonstrating that localization of the protein in VGLUT1 SVs is critical for Zn^2+^ mediated regulation of glutamatergic neurotransmission. These data emphasize coupling of transporter activity of VGLUT1 and ZnT3 leading to regulation of synaptic quantal size.

## DISCUSSION

### Corelease of multiple neurotransmitters from the same vesicle

Release of multiple neurotransmitters from the same neuron or synapse is well documented (El Mestikawy et al., 2011; Granger et al., 2017; Hnasko and Edwards, 2012). Here, we report quantitative insights on corelease of transmitters from the same vesicle and its impact on synaptic function. Using advanced super resolution imaging in combination with a robust labeling strategy and electron microscopy, we show prevalence of corelease by quantifying diverse SVs based on their transporter phenotype. Next, we validated the unexpected abundance of VGLUT1-ZnT3 SVs using quantitative biochemical and mass spectrometry analysis. Finally, using electrophysiological recordings and *in vitro* experiments, we demonstrate physiological implications of transporter colocalization on synaptic function. We show that 1) nearly ∼36% of all synaptic vesicles carry two vesicular transporters and are thus capable of storing and releasing distinct neurotransmitters, 2) however, the SV type containing VGLUT1 and ZnT3 exceedingly accounts for the majority of multitransporter vesicles (∼34% of all SVs), 3) the multitransporter SVs are more diverse than anticipated suggesting corelease of all combinations of dual neurotransmitters, 4) colocalization of VGLUT1 and ZnT3 increases vesicular glutamate content leading to an enhanced post synaptic response. We propose that, localization of distinct VTs on the same SVs may signify a specific mechanism to regulate transmitter quantal size and synaptic function. However, the physiological consequences for different VT colocalizations will depend on many factors

Since only ∼2% of all SVs (excluding those containing VGLUT1) contain more than one VT, we conclude that the majority of neuronal SVs are specific for only one neurotransmitter. However, since this quantification pertains to the whole brain, the relevance of low abundant SVs could be significant if they are enriched in specific brain regions. It is also possible that a small proportion of SVs may carry more than two distinct transporters as detected at single cell level (bar plot in Figure S1B), which were not detected in this study. Nonetheless, the unexpected combinatorial diversity of colocalized VTs reveals a new dimension of synaptic fine-tuning that extends well beyond the hitherto documented cases of cotransmission. Although the colocalization pattern of transporters discovered in this study supports, in large, existing knowledge, the data reveals some surprising combinations as well. For example, glutamate-GABA corelease has been mostly observed at synapses containing VGLUT2 and VGAT (Root et al., 2014; Shabel et al., 2014). However, our data shows that both VGLUT1 and VGLUT2 exhibit similar levels of colocalization with VGAT. This discrepancy arises, perhaps, because VGLUT1 expressing cortical synapses have not yet been extensively studied for corelease. The findings also reveal that only a small fraction of VGAT positive SVs express another transporter although increasing evidence suggests widespread incidences of cotransmission of GABA with other transmitters. This could either reflect the presence of segregated pools of SVs containing GABA and other transmitters at the reported synapses or such synapses only make up a small component of the whole inhibitory circuit.

Single vesicle investigations using whole brain preparations have been performed to address various cell biological questions (Chantranupong et al., 2020; Farsi et al., 2016; Mutch et al., 2011). For example, Mutch et al. profiled copy numbers of selected SV proteins using single vesicle imaging to assess the precision of protein sorting to vesicles. They showed little intervesicular variation in the copy numbers for SV2, VATPase, VGLUT1 and synaptotagmin 1 when compared to Syb2, Syp and Synaptogyrin indicating differential protein sorting fidelity between SV proteins (Mutch et al., 2011). However, the current study reveals a highly variable copy number for VGLUT1, suggesting the requirement for a direct evaluation of copy numbers of SV proteins in specific and intact synapses.

It should be noted that mere localization of multiple VTs on individual SVs does not imply corelease because, for some neurotransmitters, this also requires the expression of the synthesizing enzymes. On the other hand, it cannot be excluded that some VTs may allow for loading and release of multiple transmitters even in the absence of other VTs. This may be particularly relevant for the VTs known for a broader substrate specificity such as VGAT, which transports both GABA and glycine, and the VMATs, which transport all monoamines including serotonin and histamine.

Our study confirms and expands on earlier reports suggesting that the function of expressing different VTs on the same vesicle may not be confined to co-release. For instance, it was shown earlier for neurons expressing VGLUT3 together with either VAChT or VMAT2 that glutamate “synergistically” increases acetylcholine or dopamine accumulation (Gras et al., 2008; Hnasko et al., 2010) but a direct evidence of both transporters being present on the same vesicle was lacking. We report direct colocalization of two different transporters, VGLUT1 and ZnT3, in a large population of SVs and demonstrate that one of the transporters facilitates the activity of the other one leading to changes in quantal size of glutamate release. Although such a high degree of colocalization is surprising, it is consistent with the detection of ZnT3 in VGLUT1 positive excitatory synapses (McAllister and Dyck, 2017) and selective enrichment in VGLUT1 immunoisolated SVs (Gronborg et al., 2010; Salazar et al., 2005). Indeed, ZnT3 SVs containing transporters other than VGLUT1 have also been detected but they are very sparse. As stated earlier, the relevance of these low abundant SVs can only be known by region specific investigations.

Interestingly, our findings lend further support to the notion of vesicle heterogeneity at individual synapses despite the VTs following similar trafficking routes (Salazar et al., 2005). In dopaminergic neurons, glutamate and dopamine release from the same terminals is differentially regulated, suggesting differential sorting of these VTs into different SV subsets (Silm et al., 2019). However, it remains to be explored whether VGLUT1 SVs containing ZnT3 are differentially regulated from those lacking it in the same synapses, particularly in the light differences in the AZ structure between dopaminergic and glutamatergic synapses (Banerjee et al., 2021). Furthermore, the new data provide ultrastructural evidence to earlier findings of temporally coinciding release of Zn^2+^ and glutamate at specific synapses (Lavoie et al., 2011; Vergnano et al., 2014). Together, the prevalence of VGLUT1 and ZnT3 coexpressing vesicles and the influence on quantal glutamate underlies a widespread and novel regulatory mechanism of systemic synaptic excitation.

### Synaptic and circuit function implications of high overlap between VGLUT1 and ZnT3

Corelease of neurotransmitters from the same SV pools has been demonstrated in some synapses but the effect on synaptic computation and circuit function has been addressed only in a few studies (Liu et al., 2013; Shabel et al., 2014). Since the functional implications of corelease of distinct transmitters depend on various factors including the identity of neurotransmitters, the localization of pre- and postsynaptic receptors and activity-dependent changes, if any, in all of these factors, a generalized scheme of synaptic signaling is difficult to achieve. However, a common feature can be hypothesized for SVs specific for particular dual transmitters. For VGLUT1-ZnT3, the effect of the dual transmitters, glutamate and Zn^2+^, on the postsynaptic receptors is known; they both bind postsynaptic AMPA and NMDARs. Intriguingly, they exert antagonistic function, with glutamate exhibiting an excitatory response on postsynaptic neurons while Zn^2+^ inhibiting the same (Kalappa et al., 2015; Vergnano et al., 2014) indicating that Zn^2+^ might primarily act to fine tune the excitation window, as observed in glutamate-GABA coreleasing neurons (Shabel et al., 2014). It will be interesting to test if the facilitation of glutamate release by Zn^2+^ is part of a synaptic adaptation mechanism to compensate for NMDAR blockade and the resulting reduced postsynaptic calcium influx by synaptically released zinc.

Considering the high abundance of VGLUT1-ZnT3 SVs from single vesicle imaging experiments, Zn^2+^ action could underlie a systemic and robust mechanism to modulate excitation. When compared to the inhibitory circuit, mediated by GABA that predominantly balances excitation through a separate population of inhibitory neurons, Zn^2+^ is spatio-temporally locked with glutamate release to modulate excitation. Indeed, synaptic zinc, released by short trains of stimuli, selectively inhibits postsynaptic GluN2A-NMDARs and causes changes in synaptic integration and plasticity (Vergnano et al., 2014). It is, however, highly conceivable that such a direct effect depends on the magnitude of synaptic Zn^2+^ at a particular synapse, which relies on the expression level/copy number of ZnT3 in individual SVs. As the copy number of ZnT3, as determined in the present study, is highly heterogeneous (between 1-8), a region-specific zinc action depending on the expression level of ZnT3 is more plausible.

One prominent feature of the above mechanism is that Zn^2+^, unlike other neurotransmitters, is not synthesized by an enzymatic machinery but its cytoplasmic concentration is regulated by neuronal activity (Minckley et al., 2019; Sanford et al., 2019), which could lead to activity-dependent regulation of quantal glutamate mediated by ZnT3. To our knowledge, this is the first evidence of a direct role of zinc, which has been implicated in several neurodegenerative conditions and synaptic pathologies (Frederickson et al., 2005; Kawahara et al., 2020; Trist et al., 2018), in molecular mechanisms leading to changes in synaptic strength.

What will be the physiological consequence of this facilitation at the circuit level? Tzounopoulos and colleagues recently demonstrated that synaptically released zinc reduces the gain of sound-evoked responses in primary auditory cortex, A1 (Anderson et al., 2017). Although this is anticipated due to the inhibitory effect of zinc on glutamatergic receptors, the gain of principal and specific inhibitory neurons’ responses was shown to be differentially affected, requiring clarity on the mechanism of zinc action. Based on our results, one potential explanation could be that the synaptic zinc, by enhancing vesicular glutamate uptake, increases the net release of glutamate in the circuit that specifically increases feedforward inhibition, resulting in a decline in the overall gain of A1. One way to test this hypothesis would be to directly visualize glutamate release *in vivo* following expression of glutamate sensors such as iGluSnFR. As gain modulation is increasingly shown to control the dynamic range of neuronal responses to sounds and other sensory inputs (Olsen et al., 2012), earlier observed changes in synaptic zinc *in vivo* during sensory experience (Brown and Dyck, 2002; Dyck et al., 2003) suggest a profound link between activity-dependent changes in synaptic zinc and sensory processing

### Mechanism of glutamate and Zn^2+^ dual uptake

The mechanism of zinc action on glutamate uptake is yet to be investigated. However, the existing knowledge on the transport mechanism of glutamate and zinc uptake through VGLUT1 and ZnT3, respectively, could shed light on the mechanistic understanding of the dual transmitter uptake. Accumulation of most of the neurotransmitters in the vesicle lumen is regulated by a coordinated activity of vacuolar proton ATPase (V-ATPase) and VTs. The activity of V-ATPase, which is universally present in all types of vesicles independent of the neurotransmitter stored, drives protons into the lumen, generating a proton electrochemical gradient (ΔµH^+^). The VTs, which are exclusive to specific vesicle types, then use the electrochemical gradient to drive neurotransmitter transport against a concentration gradient. As different VTs use the chemical (ΔpH) and electrical (ΔΨ) components with varied efficiencies, regulation of the two components critically influences neurotransmitter uptake (Edwards, 2007). VGLUTs are thought to be predominantly driven by membrane potential (Juge et al., 2010; Juge et al., 2006), although a growing body of evidence indicates participation of ΔpH as well (Bellocchio et al., 2000; Schenck et al., 2009). While functional studies involving liposome reconstitution or single SV imaging propose VGLUT1 as a K^+^/H^+^ exchanger (Farsi et al., 2016; Preobraschenski et al., 2014), electrophysiological recordings imply an allosteric activation of VGLUT by protons from the luminal side (Eriksen et al., 2016). Indeed, a recent cryo-electron microscopy (Cryo-EM) structure of VGLUT2 elucidates a structure-based transport mechanism that implicates both ΔpH and ΔΨ (Li et al., 2020). Another characteristic feature of the VGLUTs is their biphasic dependence on chloride. While low concentration of chloride (4 mM) is required for the transport activity of VGLUTs, absence or high concentrations of chloride substantially reduce glutamate transport. The stimulatory effect of chloride has been attributed to an allosteric activation of the transporter at low concentration (Preobraschenski et al., 2018) and, at high concentrations, chloride seems to dissipate the membrane potential and thus reduces the driving force for glutamate uptake (Preobraschenski et al., 2018; Schenck et al., 2009). A direct interaction of chloride with VGLUT and thus a competition with glutamate to exert its inhibitory effect at high concentrations has been shown by many studies (Bellocchio et al., 2000; Preobraschenski et al., 2018; Schenck et al., 2009). However, arguments counter to this notion also exist (Juge et al., 2010; Juge et al., 2006). Considering the above complexity, the facilitatory effect of Zn^2+^ discovered in the current study renders additional complexity to the mechanism of glutamate transport. Zn^2+^ transport is mediated by ZnT3 which belongs to SLC30 family of Zn^2+^ transporters known for Zn^2+/^H^+^ exchange (Eide, 2006) and thus vesicular Zn^2+^ transport is expected to directly affect ΔpH. Furthermore, being a divalent cation, Zn^2+^ is poised to play a major role in the determination of luminal membrane potential (Goh et al., 2011). Indeed, a recent Cryo-EM structure of human ZnT8, a close family member of ZnT3, elucidates exchange of two Zn^2+^ ions for two protons resulting in a net increase of two positive charge in the lumen in every Zn^2+^ transport cycle (Xue et al., 2020). Thus, by lowering ΔpH and increasing ΔΨ, Zn^2+^ transport can conveniently support vesicular glutamate uptake. Finally, formation of glutamate-Zn^2+^ complex in the vesicle lumen may also reduce active glutamate amount and osmotic pressure across the membrane, which will potentially increase the upper limit of vesicular glutamate concentration (Krezel and Maret, 2016). Such a mechanism is well established for the storage of insulin in pancreatic beta-cell granules where two Zn^2+^ ions coordinate with six insulin monomers. To understand glutamate and zinc dual transport and the facilitatory effect of Zn^2+^ on glutamate uptake, rigorous investigations are required to quantify vesicular Zn^2+^ concentration and the transport mechanism of ZnT3. The availability of highly purified ZnT3 should allow functional reconstitution in liposomes or hybrid vesicles (Preobraschenski et al., 2018) to study SV Zn^2+^ transport in the future. Furthermore, vesicular Zn^2+^ uptake assay, as carried out for glutamate in the current study, can be performed to directly measure concentration of Zn^2+^ in SVs using a specific Zn^2+^ sensor. Intriguingly, the new findings show a striking similarity between regulation of substrate uptake by Zn^2+^ in synaptic vesicles and the well-known Zn^2+^ mediated insulin uptake in secretory vesicles via ZnT8 in pancreatic beta cells. Disruption of ZnT8 activity is a major cause of Type-I diabetes leading to enhanced therapeutic efforts targeting ZnT8 (Pociot and Lernmark, 2016). It has to be seen if acute disruption of ZnT3 activity underlies glutamate or metal ion dishomeostasis, which is a hallmark of major neurological diseases (Frederickson et al., 2005).

## STAR METHODS

### RESOURCE AVAILABILITY

#### Lead contact

Further information and requests for resources and reagents should be directed to and will be fulfilled by the lead contact, Sivakumar Sambandan

#### Materials availability

Reagents used in the study were of general use and from commercial sources.

#### Data and code availability

- Mass spectrometry raw data and analyses files have been deposited at www.proteomexchange.org and are publicly available as of the date of publication. Accession numbers are listed in the key resources table.
- This paper does not report original code.
- Any additional information required to reanalyze the data reported in this paper is available from the lead contact upon request.

### EXPERIMENTAL MODEL AND SUBJECT DETAILS

#### Animals

Wistar rats and ZnT3 WT and KO mice (C57BL/6 mice, RRID:MGI:3029270) were used in the study. All experimental procedures involving the use of rats and mice were carried out in accordance with national and institutional guidelines. All animals in the study, including wild type and transgenic mice, were bred at Max Planck Institute. Both, mouse and rat SVs were prepared from ∼6 weeks old animals of both sexes. Hippocampal primary culture neurons were prepared from wistar rats.

### METHOD DETAILS

#### Vesicle preparation

Isolation of synaptic vesicles from rat brain was performed as described before (Huttner et al., 1983; Takamori et al., 2006). Typically, twenty animals were used per preparation which includes multiple fractionation steps and a final control-pore glass bead chromatography (CPG) column purification step. CPG-SVs were taken from fractions containing the highest concentration of SVs for further studies. All fluorescent imaging experiments and quantitative immunoblotting were performed using freshly prepared SVs, avoiding pelleting or freezing that causes aggregation

LP2 fraction of mice SVs were prepared following a similar protocol as that of rats except that the volume of the buffers were scaled down to half because of the small size of mouse brains. Typically, 10 wildtype (WT) mice and 10 knockout (KO) mice were sacrificed together and the SV isolation for WT and KO was performed simultaneously.

Chromaffin granules were isolated from bovine adrenal medulla by combining differential and discontinuous (0.3 M/1.8 M) sucrose gradient centrifugation (Birinci et al., 2020).

#### Dynamic Light Scattering

Dynamic light scattering (DLS) was used to determine the size distribution and integrity of synaptic vesicles following purification. Measurements were carried out with 25 µl of SV sample, using a DynaPro™ Dynamic Light Scattering Instrument (Wyatt Technology) at a laser power of 5 % with 20 s acquisition duration. Individual measurements were performed 10 times and the values were averaged.

#### Electron microscopy

To evaluate the purity and quality of prepared synaptic vesicles, negative staining was regularly performed. The sample was bound to carbon coated, glow discharged grids and counterstained using 1% uranyl acetate. For immunogold-labeling (IEM, Figure S4), purified SVs were absorbed to formvar coated grids, fixed with 4% paraformaldehyde, quenched with 20 mM glycine and immunostained using the described antibody, followed by addition of Protein A-gold (10 nm). The grids were then rinsed repeatedly with TPBS and high-salt TPBS (0.5 M NaCl) and post-fixed with 2% glutaraldehyde. For double labeling (Figure 3), the first immunoreaction was blocked by fixation with 1% glutaraldehyde, followed by a second immunogold labeling. After counterstained with 1% uranyl acetate, samples were investigated using Talos L120C (Thermo Fisher Scientifc). The antibodies were used in reverse order as well to exclude the effects of glutaraldehyde fixation on the antigenicity. Antibodies used in IEM are published previously (Syp, clone G95, rabbit polyclonal, (Jahn et al., 1985)) and Synaptic Systems, Germany (Syb2-104211; VGLUT1-135303; ZnT3-197011.

#### SV immunolabeling for DyMIN STED Imaging

Purified SVs were resuspended in blocking solution [PBS, pH 7.4, 5% normal goat serum (v/v)] with constant rotation for 1 h at room temperature (RT). Primary antibodies or fluorophore conjugated primary antibodies were added in blocking solution containing SVs at a concentration of 6µg/ml. with constant rotation for 1 h at RT. Subsequently, the SV sample was subjected to size exclusion chromatography (SEC) using a superdex 200 column by means of ÄKTA system (GE Healthcare) to purify labeled SVs from unbound antibodies. As expected, the SVs were eluted in void volume and the antibodies and excess serum proteins were eluted via the stationary phase, resulting in clear separate peaks for SVs and smaller contaminants (Figure S2). The SVs were typically eluted at∼14ml elution volume corresponding to elution fractions 36 to 37.

Following SEC purification, SV integrity was again analyzed by DLS. In some cases, the SV and antibody peaks were also analyzed by EM and DyMIN STED. The SV peak displayed small round profiles in the EM typical of synaptic vesicles, which was lacking in the antibody peak (Figure S2). DyMIN STED, as expected, detected fluorescence puncta in both peaks (Figure S2). DLS count was taken as a substitute for SV concentration, which allowed us to use the same amount of SVs (1000 counts/s) for immobilization on glass bottom dishes (Mat Tek, P35G-1.5-14-C) by incubating at 4°C for 1 h. Afterwards, SVs were fixed using 4% paraformaldehyde (Thermo scientific, 28908) in PBS (v/v) for 5 min and washed thrice in PBS at an interval of 5 minutes. If secondary antibody labeling was needed, the immobilized SVs were incubated in blocking solution for 30 minutes followed by secondary antibody incubation at a concentration of 8µg/ml (a dilution of 1:250) for 1 h at RT. Afterwards, samples were washed thrice with PBS at an interval of 5 minutes. Finally, mounting media (Invitrogen P36961) was applied on top of SVs and images were typically acquired within 48 h of sample preparation.

For dual color imaging in purified SVs, both mouse and rabbit antibodies of vesicular transporters were used except for VMAT2 and VAChT, for which only rabbit antibodies were used. To determine the proportion of SVs containing a specific transporter (Figure 2D), rabbit or mouse Syp antibody was used. Swapping the antibody species between Syp and respective transporters did not change the outcome. In some experiments where usage of the same species antibodies were required, secondary nanobodies were pre-mixed with primary antibodies and used (multiplexing). VMAT2 and VAChT staining using multiplexing did not result in reproducible results and was not included in the analysis. The total number of analyzed vesicles (n) in the colocalization experiments (Fig 2C-J) was more than 20,000 (20K) for VGLUT1, n>12K for VGLUT2, n>3K for VGLUT3, n>10K for VGAT, n>3K for VMAT2, n>3K for VAChT and n>20K for ZnT3.

#### DyMIN STED imaging

DyMIN STED nanoscopy was performed using a quad scanning STED microscope (Abberior Instruments, Göttingen, Germany) equipped with a UPlanSApo 100x/1.40 Oil immersion objective (Olympus, Tokyo, Japan) and the optical set-up was described earlier (Heine et al., 2017). The pinhole was set to 1.0 Airy units and a pixel size of 15 nm was used. In dual color DyMIN STED imaging, Abberior Star Red (Ab.St.RED) and Alexa Flour 594 (AF594) combination was used while Alexa Fluor 488 (AF488), if required, was used as a reference marker in confocal mode. Ab.St.RED was excited at 640 nm and AlexaFluor 594 was excited at 561 nm while STED was performed at 775 nm wavelength. The fluorescence signal was detected using avalanche photo diodes with bandpass filters and a gating of 0.75–8 ns was applied. Pixel dwell times of 10-20 µs were used. Each line was scanned 3 times and the signal was accumulated. The typical image size for purified SVs was 50 µm^2^. The number of images per transporter pair ranged from 10 to 50 per experiment. In comparisons between DyMIN and conventional STED and confocal acquisitions shown in Figure 1, the same excitation power for confocal and STED was used. The STED power in conventional STED corresponded to *P*_max_ of DyMIN STED (final step).

#### Colocalization analysis

To determine positive vesicular colocalization in DyMIN STED fluorescence images, we tagged epitope pairs that are known to be colocalized, using two spectrally distinct dyes and calculated the nearest neighbor distance (NND) of all the puncta in one channel with respect to the other. As expected, the distribution of the NND values followed a Rayleigh distribution, as described earlier (Geumann et al., 2008). For example, we labeled two different epitopes of the same protein, Syp, with two distinct colors using a mouse (clone 72.1) and a rabbit Syp (clone G95) antibodies and calculated NND (Figure S4E-G). Although the corresponding images looked like mirror images of each other, we obtained a NND distribution with a median of approximately 15.

Next, we performed a similar task but targeting two different proteins, Syp and Syb2, that was shown to be expressed in all SVs (Figure S4I-J) and again calculated NND for all spots. Since Syp and Syb2 are the two most abundant proteins by copy number (Wilhelm et al., 2014), we supposed that the fluorescent spots would be larger for both channels and thus yielding a larger NND distribution (median ∼25 nm). The above two positive controls provided a realistic range for inter spot distances when the proteins of interest are closest (NND <30 nm; two epitopes on the same protein) or farthest (NND <50 nm; Syp and Syb2) on the same vesicle. From the above experiment, we adopted that that any two proteins will be considered as colocalized on the same vesicle if the NND between the coordinates is less than 50 nm.

For colocalization analysis, the dual color images are median filtered using a radius of one pixel (15 nm) to remove single pixel background. Then, the coordinates of individual SVs in the two images were extracted simultaneously using an open source plug-in for Fiji, ThunderSTORM (Ovesny et al., 2014). Using a custom written R script, the distance *d* between two puncta with coordinates (*x*_1_, *y*_1_) and (*x*_2_, *y*_2_) in the paired images was calculated using the formula,

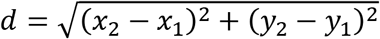

For colocalization analysis in culture neurons (Figure S6E), images were background subtracted using Rolling ball method with a radius of 50 pixels and colocalization correlation was performed using Coloc 2 plugin in Fiji. Costes’ test (p=1) was repeated 100 times to test statistical significance of the determined colocalization coefficients

#### Neuronal Cell Culture and Immunolabeling

Cultures of hippocampal neurons were prepared from Wistar rats at postnatal day P0–P1 using standard procedures (Kaech and Banker, 2006). For immunostainings, cells (16-22 DIV) were washed with PBS and fixed in 4% PFA in PBS (pH 7.4) for 20 min at RT, quenched with ammonium chloride and glycine (100 mM each) for 5 min, permeabilized with 0.1% Triton X-100 for another 5 min, and blocked in PBS supplemented with 1% BSA for 30 min. Both the primary and secondary antibody incubations were performed in PBS for 1 h at room temperature. Samples were mounted in ProLong Diamond Antifade Mountant medium (Thermo Fisher, P36961).

### MINSTED DNA PAINT

Cultured hippocampal neurons, fixed (4% PFA for 10 min), quenched (100 mM NH4Cl2 for 10 min), permeabilized (5% TritonX-100 for 5 min) and blocked (5% NGS for 1 h), was indirectly immunolabeled against VGLUT1, ZnT3 and Piccolo (135308, 197011, SySy, 1:200 dilution) and Piccolo (142104, SySy, 1:500 dilution). The anti-mouse and anti-rabbit secondary antibodies were coupled to P1 and P3 docking strands at the 5’ end (5’-3’, P1 - TTA TAC ATC TA; P3 - TTT CTT CAT TA; Massive Photonics, 1:100 dilution). The imager strands contained Cy3B at the 3’ end (5’- 3’, P1-CTA GAT GTA T; P3 - GTA ATG AAG A),) and the anti-guinea pig antibody was coupled to AF488 (1:500 dilution). The sample was post-fixed using 4% PFA following a washing step.

MINSTED nanoscopy enables an enhanced resolution by localizing single emitters with an effective STED PSF of FWHM d (Weber et al., 2021). Thus, the information from all emitted photons N can be used in order to find the emitter’s position to an uncertainty σ scaling with 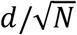, leading to further downsizing of STED resolution to the molecular scale. Here, localizations were performed at *d* = 30 nm which led to a median localization precision below 1.5 nm within a maximum duration of 200 ms per localization. MINSTED measurements were carried out in PBS buffer containing the respective imager strand (5 nM) and MgCl_2_ (75 mM). Initially, the piccolo signal (AF488) was acquired in confocal mode in a large field of view and a synaptic region of interest was selected for two-color MINSTED imaging. Multiplexing was performed by sequential image acquisition with two different imager strand solutions containing P1 or P3 sequences (Schnitzbauer et al., 2017).

For image analysis, the localizations from two-color MINSTED images were filtered (27% of the localization events were discarded) and were subsequently clustered. Cluster analysis was performed by initially segmenting regions of high localization density using an in-house developed tessellation algorithm as performed elsewhere (Levet et al., 2015). The triangles of a Delaunay triangulation of all localizations 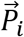 were assigned to segments if the triangles’ corners were attributed a localization density *D*_*i*_ greater than the average localization density in the image and the triangles’ edges were shorter than 20 nm. The localization density *D*_*i*_ at the corner 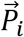 was estimated by *C*_*i*_/*A*_*i*_, where *C*_*i*_ is the number of unique corners and *A*_*i*_ the combined area of the triangles attached to 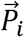. Non-assigned localizations were subsequently added to their closest segment if not further away than 30 nm with respect to the segment’s boundary. After discarding those segments with ≤ 10 localizations, a total of 863 and 504 clusters were found for two-color- and VGLUT1-clustering, respectively. For two-color clustering, all localizations of both colors were merged and the clustering was performed “color blind”. Identifying the colors of the assigned localizations after clustering enabled to quantify VGLUT1-ZnT3-colocalization for each cluster. Based on the same dataset, VGLUT1-clustering was performed only on the data from the VGLUT1 color channel. In total, 22 two-color MINSTED images and 59 AZs from four different experiments were analyzed.

#### Copy number measurement

SVs were labeled for the protein of interest using monoclonal primary antibody (VGLUT1 – synaptic systems, Cat. No. 135011; ZnT3 synaptic systems, Cat. No. 197011) and secondary nanobody directed against the kappa light chain of mouse IgG (FluoTag®-X2, NanoTag Biotechnologies, Germany, Cat. No. N1202). Two fluorophores (Star RED, Abberior) are site-specifically coupled to one nanobody molecule ensuring a fixed stoichiometry of 4 to 1 (two nanobodies bind to each F(ab) of IgG molecule) for the number of fluorophores per IgG (Shaw et al., 2019; Sograte-Idrissi et al., 2020). Initially, 2.5-fold excess of secondary nanobody was added to the monoclonal primary antibody to saturate available binding sites of IgG. Then, the antibody-nanobody mixture was diluted 5 times using PBS and incubated for 1h at RT with subtle shaking. The pre-incubated antibody-nanobody complex was diluted using blocking solution to reach concentrations optimized for the primary antibody and incubated with SVs with constant rotation for 1h at RT. The labeled SVs were purified using SEC as described earlier, followed by immobilization and imaging. For the calculation of copy number, the fluorescence intensity of each vesicle puncta was divided by the mean fluorescence intensity of puncta corresponding to free antibody-nanobody complexes.

#### Immunoisolation

Immunoisolation of SVs was performed using paramagnetic Dynabeads (Takamori et al., 2000; Zander et al., 2010). 50 µl of Protein G Dynabead slurry was used per reaction. After washing the beads with 1 ml PBS (pH 7.4) three times with an interval of 10 min, ZnT3 antibody was added at a concentration of 24 µg/ml and incubated for 1hr at 4°C with constant rotation. The antibody-beads mixture was washed once with PBS for 15minutes. This mixture was then incubated with 50 µg of LS1 protein overnight at 4°C under constant rotation. Next day, LS1-protein-dynabead mixture was washed thrice with an interval of 15 min each, using 1xPBS supplemented with protease inhibitors. Beads were transferred into new low binding Eppendorf tubes and spun down, placed on a magnet to completely remove the buffer from the beads. Afterwards, 50 µl of 5xLSB containing 10% Beta-mercaptoethanol was added to the bead pellet and the sample was subjected to heating at 70°C for 10 min with 750-800 rpm stirring. The vesicle concentration curve was run by increasing protein amount of LS1 with respect to IPs of ZnT3 and Syb2 (Schagger, 2006).

#### Quantitative immunoblotting

Determination of the amount of ZnT3 in SVs was performed using quantitative western blotting of SV proteins (CPG-SVs) in comparison to a standard curve of a highly purified full-length recombinant protein (Takamori et al., 2006). Increasing amount of SV and recombinant protein samples were loaded in the same gel and all measurements within the linear range of the standard curve were taken into account. Protein bands were visualized using an Odyssey CLx scanner (Li-cor) and signal intensities were analyzed using Fiii.

#### Glutamate uptake assay

LP2 fractions of SVs were prepared according to the protocol described previously (Preobraschenski et al., 2014) from adult (5-8 weeks old) WT and ZnT3 KO mice brains. WT and KO LP2 samples for all glutamate uptake assays were prepared together to avoid differences in specimen quality. 15 µg of SVs was added to 90 µl of uptake buffer (300 mM Glycine, 10 mM MOPS-KOH, 2 mM MgSO4, pH 7.3) and incubated with 10 µl of 10X ATP-glutamate mix (10 mM potassium glutamate, 4 mM MgATP, 4 mM KCl) in a microtiter plate at 37°C for 15 minutes. 10X ATP-glutamate mix was adjusted accordingly for the uptake assays in the presence of Zn^2+^. After 15 min, 90 µl of sample was transferred in a microfilter plate (MultiScreenHTS + HiFlow FB Filter Plate, Merck Millipore) and filtered using vacuum manifold while being washed thrice with 200 µl ice cold uptake buffer. In order to rupture SVs and measure fluorescence change, 50 µl iGluSnFR solution (1µM purified iGluSnFR protein, 0.1% Triton X-100 resuspended in uptake buffer) was added per well and incubated for 30 min with gentle shaking at RT, protected from light. Fluorescence was measured using microtiter plate reader at excitation and emission wavelengths of 495 nm and 520 nm, respectively. Glutamate uptake in the presence of 30 µM FCCP was used as the baseline fluorescence and subtracted from all measured values. Solutions with different Zn^2+^ concentrations were prepared using ZnSO_4_ in different metal buffering systems (Qin et al., 2011; Vinkenborg et al., 2009). 1 pM of Zn^2+^ was used as the basal level of zinc (control) in uptake assays. For obtaining iGluSnFR standard curve, different glutamate concentrations ranging from 0 to 10 mM was added to uptake buffer (300mM Glycine, 10mM MOPS-KOH, 2mM MgSO4, pH 7.3) to make a final volume of 100 µl. The mixture was incubated in a microtitre plate at 37°C for 15 minutes. Afterwards, 50 µl iGluSnFR solution (1µM purified iGluSnFR protein, 0.1 % Triton X-100 resuspended in uptake buffer) was added per well and incubated for 30 min with gentle shaking at RT, protected from light. Later, the flourescence was measured using a microtitre plate reader at excitation and emission wavelengths of 495 nm and 520 nm respectively. Plasmid containing His6-tagged iGluSnFR variant, SF-iGluSnFR.A184S (Marvin et al., 2018), was kindly provided by L. Looger and purified by Ni-NTA affinity chromatography (Marvin et al., 2013).

#### Hippocampal slice preparation

WT (C57BL6/J) and ZnT3 KO mice (P19–P27, either sex) were anesthetized by diethylether and decapitated immediately. The brains were quickly removed, and horizontally sliced (400 μm) in ice-cold cutting buffer using a VT1200S vibratome (Leica). The cutting buffer composition (composition in mM): 26 NaHCO_3_, 1.25 NaH_2_PO_4_, 4 KCl, 10 glucose, 230 sucrose, 0.5 CaCl_2_, and 10 MgSO_4_ (pH 7.4, osmolarity ∼305 mOsm/L; equilibrated with 95% O2 and 5% CO_2_). The brain slices were then transferred to a custom-msade holding chamber containing the ACSF solution and incubated at 37°C for at least 30 min, and further recovered at room temperature for more than 1 h before recording. ACSF solution (composition in mM): 125 NaCl, 26 NaHCO_3_, 2.5 KCl, 1.25 NaH_2_PO_4_, 1 MgCl_2_, 2 CaCl_2_, and 25 glucose (pH 7.4, osmolarity ∼305 mOsm/L; equilibrated with 95% O_2_ and 5% CO_2_). All experiments complied with national animal care guidelines, and were approved by the University Medical Center Göttingen board for animal welfare and the animal welfare office of the state of Lower Saxony.

#### Electrophysiology

Glass patch pipettes were pulled from borosilicate glass using a GB150TF-10 (Science Products) and had resistance of 2–5 MΩ. Brain slices were placed in the custom-made recording chamber kept at 37°C and were continually perfused with oxygenated ACSF (rate: 2.5ml/min). 10 μM bicuculine methiodide, 50 μM DAP-5 and 1μM tetrodotoxin (TTX) were added to ACSF during the recordings. Cells were examined in a whole cell configuration under voltage clamp mode at a holding potential of −70 mV. Recordings were made using a custom-assembled rig including a microscope (Zeiss), a Patchstar 360 micromanipulator (Scientifica), a PPS2 perfusion system (Multichannel system) in a Faraday cage, and a HEKA EPC10 amplifier. Data were acquired with Patchmaster software (HEKA) at an acquisition rate of 20 kHz, and low-pass filtered at 2 kHz. During recordings, glass pipettes were filled with an internal solution (composition in mM): 130 CsMeSO_4_, 5 TEA-Cl, 5 NaCl, 10 HEPES, 4 MgCl_2_, 0.1 EGTA, 10 Na-creatinephosphate, 4 ATP and 0.4 GTP, pH 7.35, osmolarity ∼300 mOsm/L. Zn^2+^ application is achieved by 5 min perfusion of ACSF containing 200 μM ZnCl2. Uncompensated series resistance (Rs) was monitored by the delivery of 5 mV voltage step at defined points during the recording. Recordings were stopped or excluded if the Rs exceeded 20 MΩ, or changed by more than 20%. Events were analyzed by a custom-written macro in Igor Pro 6.36 software, as previously described (Clements and Bekkers, 1997).

#### Single cell transcriptome analysis

Raw expression values for each gene in each cell were normalized to the highest expression gene of the eight transporters to compare them across all cells. The gene expression levels are assigned between the value 0 and 1, where 1 means highest expression of the gene in that cell. If two genes had exactly the same expression in the same cell, both of these were assigned a value of 1. We clustered the co-expressed genes inside different cell types to generate a heat map. For cluster analysis, we used the Euclidean method as a clustering distance and default hierarchical clustering method in complex Heatmap R package version 2.5.4. To define cell types, we considered a gene expressed in a cell type if its normalized expression is more than 0.5 (meaning that its expression is more than half of the highest expressed in that cell type). After obtaining the expression of unique genes expressed in each cell type, we quantified cell types that exhibited expression of more than one of the eight transporter genes.

#### Sample preparation for mass spectrometry

Purified synaptic vesicles were lysed using an equal volume of the lysis buffer (4% (wt/v) SDS, 100 mM HEPES, 1 mM EDTA in water), sonicated for 10 min using 30 s on/30 s off cycles at the maximum output of Bioruptor ultrasonication device (Diagenode, Seraing, Belgium). Protein concentration was assessed using BCA assay kit (Thermo Fisher Scientific, Bleiswijk, Netherlands) according to manufacturer’s instructions. Cysteine moieties were reduced and alkylated using 10 mM tris(2-carboxyethyl)phosphine (TCEP) and 40 mM chloroacetamide (CAA) at 37 °C for 30 min. UPS2 standard (Sigma-Aldrich, Taufkirchen, Germany) was reconstituted in the lysis buffer and cysteines were reduced and alkylated as described above. 1.5 µg of UPS2 proteins were spiked into 9 µg of SV sample. Additionally, 1.5 µg of UPS2 standard without SV proteins and 9 µg of SV proteins without UPS2 standard were prepared. Proteins were purified using magnetic beads with carboxylate modified hydrophilic and hydrophobic surface (GE Healthcare, Buckinghamshire, UK) as described in (Hughes et al., 2019). Proteins were pre-digested in 50 mM ammonium bicarbonate buffer for 2 h at 37°C using recombinant LysC-protease (Promega, Madison, USA) at 1:100 LysC-to protein wt/wt ratio followed by the overnight digestion at 37°C using MS-grade trypsin (Promega, Madison, USA) at 1:20 trypsin-to-protein wt/wt ratio. Peptides were dried in a centrifugal Savant SpeedVac vacuum concentrator (Thermo Fisher Scientific, Waltham, USA), reconstituted in 2% (v/v) acetonitrile (ACN), 0.1% (v/v) trifluoroacetic acid (TFA) in water and subjected to LC-MS/MS analysis.

#### LC-MS/MS

Digested protein samples were analyzed in technical (injection) duplicates (SV samples with or without UPS2 proteins) or as a single injection (UPS2 proteins only) on a hybrid quadrupole-ion trap-orbitrap instrument (Orbitrap Fusion, Thermo Fisher Scientific, San Jose, USA) using data-dependent (DDA) and data independent (DIA) acquisition methods. Peptides were concentrated onto a C18 PepMap100-trapping column (0.3 mm x 5 mm, 5 µm, Thermo Fisher Scientific, Waltham, USA) connected to an in-house packed C18 analytical column (75 µm x 300 mm; Reprosil-Pur 120 C18-AQ, 1.9 µm, Dr. Maisch GmbH, Ammerbuch, Germany). Liquid chromatography was operated on an UltiMate-3000 UHPLC nanosystem (Thermo Fisher Scientific, Waltham, USA). The columns were pre-equilibrated using a mixture of 95% buffer A (0.1% (v/v) formic acid (FA) in water) 5% buffer B (80% (v/v) ACN, 0.1% (v/v) FA in water). Peptides were eluted using a 118 min linear gradient from 5 to 7% buffer B over 3 min, from 7 to 20% over the next 57 min, from 20 to 32% over 30 min, and from 32 to 50% over 16 min. Elution was followed by a washing step at 90% buffer B over 6 min and a re-equilibration step at 5% buffer B over 6 min. MS was operated in DDA mode with the following settings: MS1 scans of 350-1650 *m/z* range were acquired in positive mode at the resolution of 120000 at 200 *m/z*, 300% automated gain control target (AGC), and 50 ms maximum injection time (IT). Most abundant peptide ions (charge states 2-7) were selected for MS/MS using an isolation window of 1.6 *m/z*. Peptide fragments were generated in an HCD cell at 28% normalized collision energy (NCE) and measured in Orbitrap at the resolution of 15000 at 200 *m/z*, normalized AGC of 1000%, and maximum IT of 54 ms. The duty cycle was kept at 2.5 s and dynamic exclusion at 30 s. When operated in DIA mode, MS1 spectra were acquired as in the DDA mode. Each MS1 scan was followed by 40 MS/MS scans of variable isolation window width. Peptide fragments were generated in an HCD cell at 30% NCE and measured in Orbitrap at the resolution of 30000 at 200 *m/z* and maximum IT of 54 ms.

#### LC-MS/MS data analysis

Raw MS data were analyzed using Spectronaut software ((Bruderer et al., 2015), v. 14.3.200701.47784). For each biological replicate, (2 in total), a separate spectral library was built that included all DDA and DIA data from a given SV-replicate and the pure UPS2 standard protein. Spectral libraries were generated using a built-in Pulsar search engine under default settings. Specifically, *Rattus norvegicus* protein sequence database containing 29951 entries and retrieved from Uniprot (UniProt, 2019) in February 2019 was used in conjunction with UPS2 protein sequences (48 entries). Carbamidomethylation of cysteines was set as a fixed modification, and N-Terminal protein acetylation and oxidation of methionine as variable modifications. DIA-based quantification was performed separately for each biological replicate using the respective spectral library. The settings were kept default except “Q value sparse” with no imputation and no cross-run normalization were selected for data filtering. IBAQ values (Schwanhausser et al., 2011) were reported as protein group intensity values. The data was further analyzed using a custom R script. In brief, only proteins quantified with at least two peptides were retained in the data set. Decoy sequences, keratins, immunoglobulins, and trypsin were removed from the data. IBAQ intensities from the two technical replicates were averaged. Background-level intensities of proteins present in the UPS2 standard but identified in the SV sample without spiked-in UPS2 proteins were subtracted from UPS2 protein intensities in the SV sample containing spiked-in UPS2 standard. Additionally, iBAQ intensities were corrected by the protein sequence coverage and the protein concentrations were assessed using a linear model that was derived from the known protein amount of UPS2 proteins *vs.* IBAQ intensities in the first technical replicate (R^2^ = 0.614). Only proteins quantified in both biological replicates are reported. If not specified otherwise, uncertainty in the protein concentration estimation is given as 68% confidence interval based on the fitted linear model. All raw mass spectrometry and software analysis files were deposited to the ProteomeXchange Consortium (www.proteomexchange.org) via the PRIDE (Perez-Riverol et al., 2019) partner repository. The variable window setup for DIA acquisition is provided in the Supplemental Information (SI).

## SUPPLEMENTAL TABLES

Table S1: Compilation of studies on cotransmission and corelease, related to Figure 3 and S1

Table S2: SV Proteome_Quantitative mass spectrometry data, related to Figure 4

## QUANTIFICATION AND STATISTICAL ANALYSIS

For single vesicle colocalization between two images, an ‘object based’ analysis was performed, implemented in R. Thus, the colocalization quantification is based on the actual number vesicles and not pixel correlation between the two channels analyzed. The method does not assume that the fluorescent intensity profiles between the two channels are comparable, which is appropriate for single vesicle analysis. Thus, composite or merged pseudo color images, as regularly used for ‘pixel based’ analysis, were avoided in the related figures (Figure 1, 2, S5 and S6). For colocalization analysis of confocal images in culture neurons (Figure S6E), a pixel based colocalization correlation was performed using Coloc2 plugin in Fiji. Costes’ test (p=1) was repeated 100 times to test statistical significance of the determined colocalization coefficients. Statistical analysis for all experiments was performed using OriginPro 2020. In all experiments, statistical significance between two different samples was performed using unpaired t test following confirmation of normal distribution of data sets (Figure 1B-E, 4F, 7D). This includes comparison of microscopy techniques in single vesicle imaging (Figure 1B and Cs), comparison of conventional and ‘in-solution’ labeling of purified vesicles (Figure 1D and E), comparison of copy numbers of VGLUT1 and ZnT3 samples (Figure 4F) and electrophysiological experiments (Figure 7D). The statistical significance of glutamate uptakes (nine samples) in the presence of different zinc concentrations was performed using two-way ANOVA (Figure 6B). The quantification in Figure 3 is based on the colocalization degree of each VT pairs and the respective proportion of each VT in the entire total SV population shown in Figure 2. Samples sizes of vesicles in the colocalization analysis is provided in both, STAR Methods and Figure legends.

## ACKNOWLEDGEMENTS

We thank members of RJ lab for useful discussions, Raza-Ur Rahman (Broad Institute) for RSeq analysis, Brigitte Barg-Kues for genotyping and Sigrid Schmidt for preparing culture neurons. We thank Gudrun Ahnert-Hilger (Berlin), Silvio Rizzoli (Göttingen), Pablo Castillo (New York) and Agata Witkowska (Berlin) for helpful comments on the manuscript. This research project was conducted with support from Volkswagen Foundation (‘Experiment’ grant to SS), European Research Council Advanced Grant (SVNeuroTrans to RJ), the John Black Charitable Foundation (to IM), and Deutsche Forschungsgemeinschaft (SFB1286 to HU). HE is part of the Max Planck School of Photonics supported by BMBF, Max Planck Society, and Fraunhofer Society.

## AUTHOR CONTRIBUTIONS

SS conceptualized the study, performed and supervised experiments, evaluated all data and wrote the paper with inputs from all co-authors. RJ provided input and appraised all data. SWH provided input on STED and MINSTED experiments. NU, VM, JP, MG, LB, and EZ designed and performed different aspects of the study. HE performed and analyzed MINSTED experiments and ML contributed in the cluster analysis. JJ and IM performed and analyzed electrophysiology experiments. MN, IS and HU performed mass spectrometry studies and analyzed data. AP drafted and wrote the imaging analysis workflow. DR performed electron microscopy and analyzed data.

## SUPPLEMENTAL INFORMATION

Supplemental Figures

Figure S1, related to Introduction and Figure 3

Figure S2, related to Figure 1

Figure S3, related to Figure 1, 2

Figure S4, related to Figure 1, 2

Figure S5, related to Figure 2

Figure S6, related to Figure 2, 3, 5

Figure S7, related to Figure 6

**Figure S1.**
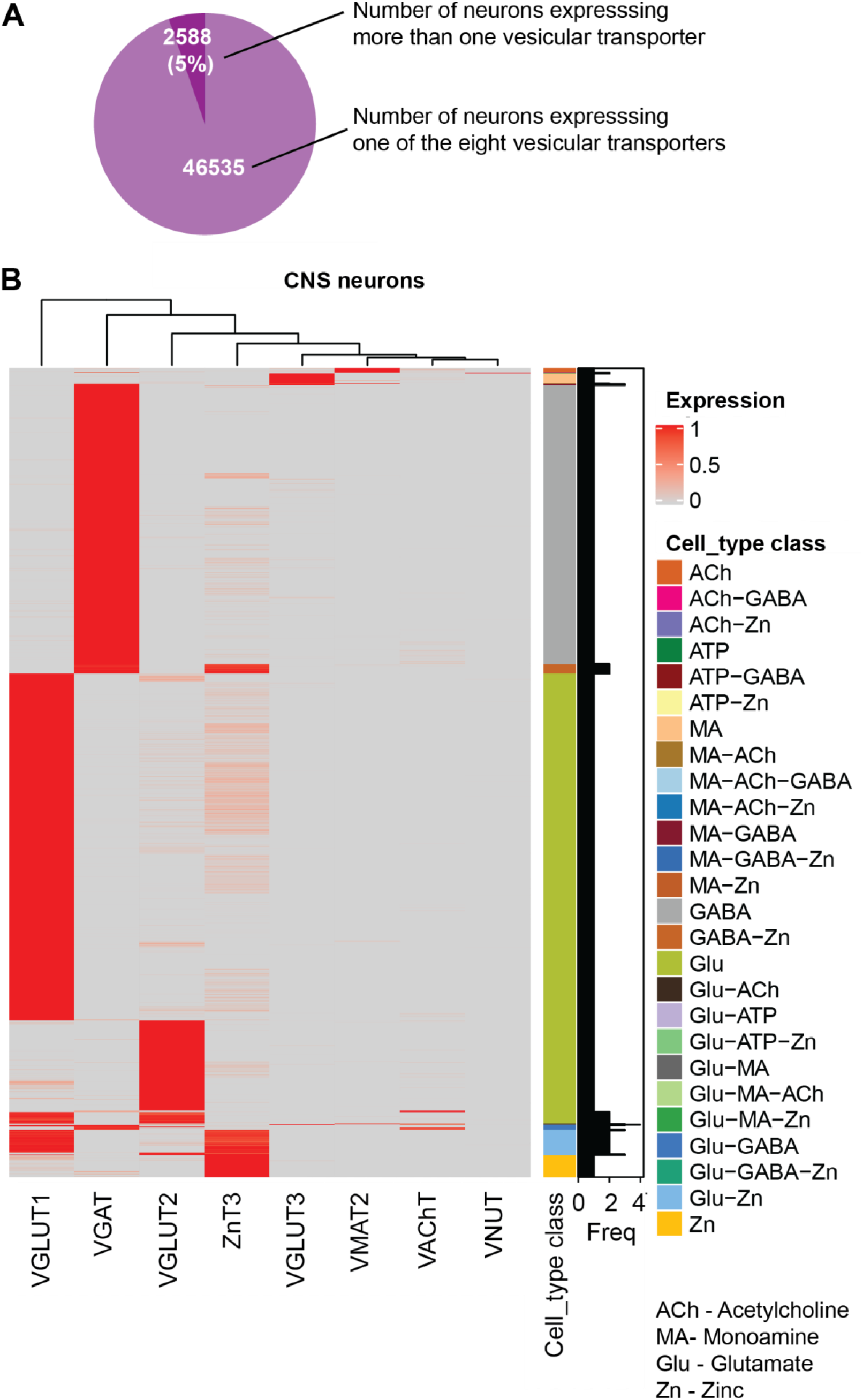
Coexpression of vesicular transporters in the whole brain revealed by single cell transcriptome analysis. (**A**) Pie diagram showing the composition of neurons in the whole brain expressing more than one vesicular transporter (total neurons analyzed = 70968; neurons expressing at least one transporter = 49123) (**B**) Heat map showing the normalized co-expression of different vesicular transporter genes. Horizontal lines represent single neurons expressing either one or multiple vesicular transporters shown on the X-axis. The heat map color ranges from grey to red (0 to 1); grey indicates no expression and red intensity shows the expression level of genes. The color bar on the right shows the assigned cell types based on the expression of one or more types of transporters. Bar plot (black) represents the number of transporter genes expressed in the cell type on the left. Note that cell types expressing three or even four different transporters are also identified. The transcriptome data is reanalyzed from Zeisel et al., 2018.

**Figure S2.**
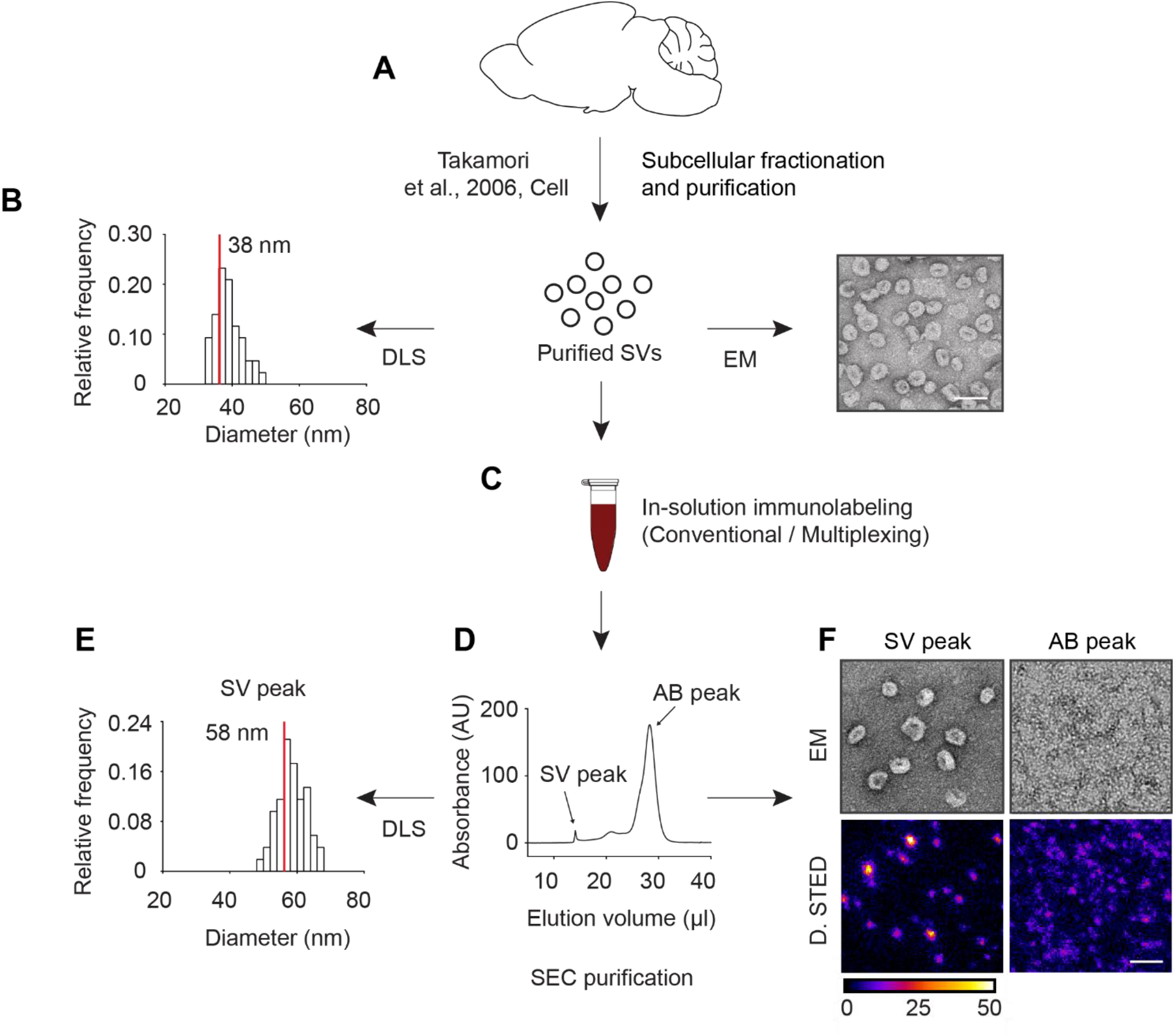
Overview of the workflow of synaptic vesicle isolation, labeling and further purification for DyMIN STED single vesicle imaging. (**A**) Purified SVs were prepared by a multi-step subcellular fractionation, as described in Takamori et al., 2006. (**B**) The purity and homogeneity of isolated SVs was tested using electron microscopy (EM, representative image on the right) and diffraction light scattering (DLS, summary histogram of particle diameter on the left). The mean size of individual SVs, as measured by DLS particle diameter was ∼38 nm (n = 22 experiments). DLS particle count was used as a measure for the concentration of individual vesicles in the further steps. Scale bar, 50 nm. (**C**) After a blocking step, isolated SVs were incubated with saturating antibody concentration ‘in-solution’ under constant rotation for 1 h. (**D**) The immunolabeled SVs were then subjected to size exclusion chromatography (SEC) to remove unbound free antibodies and other contaminants such as excess serum proteins in the blocking buffer. Representative chromatogram demonstrates clear separation between the labeled SVs (SV peak) that was eluted in void volume and the contaminants (AB peak). (**E**) The quality of SEC purification was verified by subjecting the SV and AB peaks for DLS measurements in all experiments. Note that the mean size of SVs (red line in the DLS histogram) is increased (∼20 nm) following immunolabeling indicating the change in size of SVs due to the bound antibodies. DLS measurement was not possible in the AB peak because of the heterogeneous particle size. (**F**) In some experiments, EM and DyMIN STED were used to examine SV and AB peaks following direct immunolabeling of SVs. As expected, only the SV peak contained detectable SVs in the EM, whereas fluorescence was detected in both the samples. The labeled SVs were then immobilized on glass coverslips to prepare for DyMIN STED. Scale bar, 100nm

**Figure S3.**
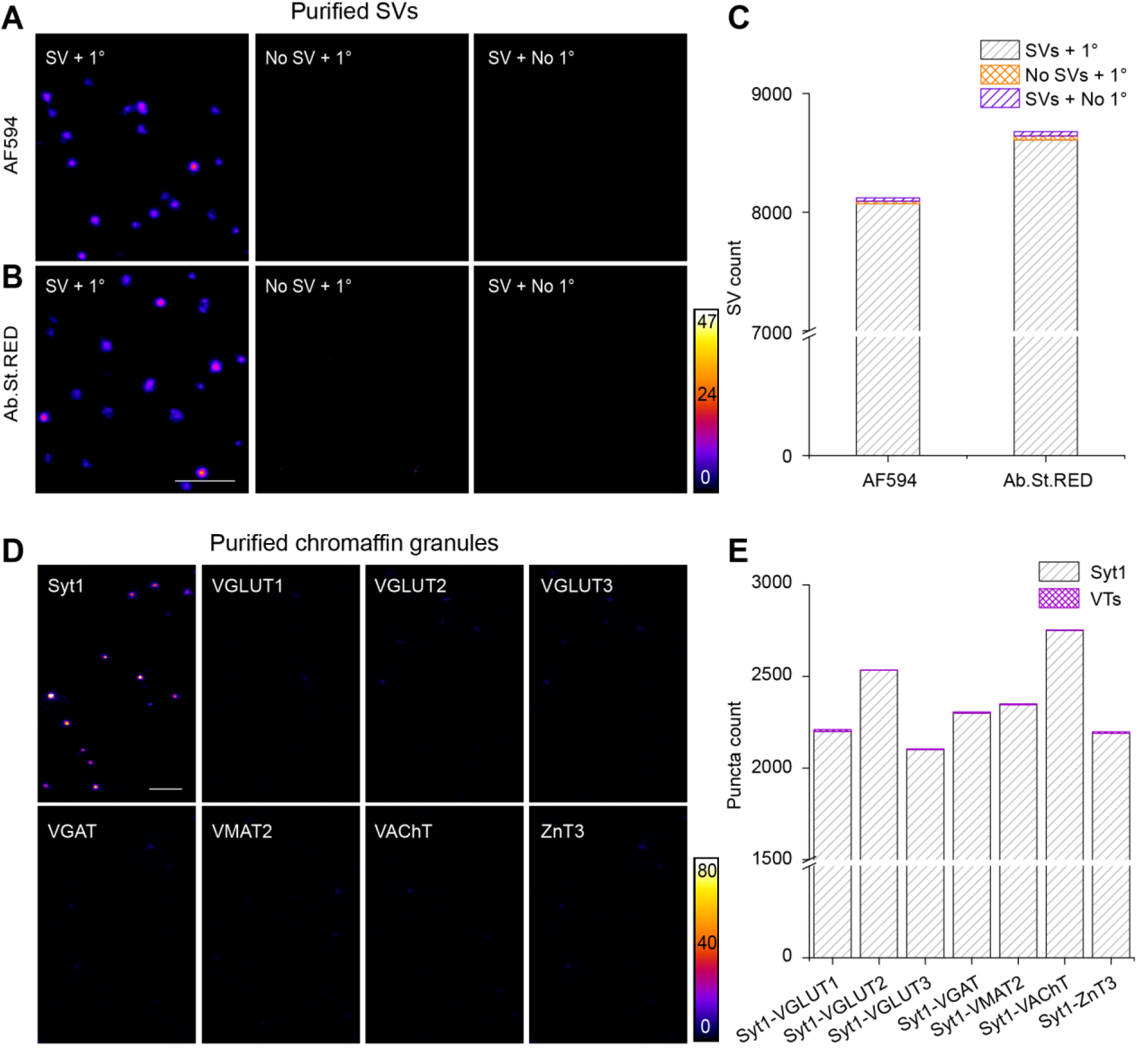
Marginal background and unspecific fluorescence in DyMIN STED single vesicle imaging. (**A**) Left, representative DyMIN STED images (fire) of purified SVs labeled against Syb2 using mouse anti-Syb2 primary antibody and anti-mouse secondary antibody conjugated with Alexa Fluor 594 (AF594). Middle, image showing undetectable fluorescence when SVs were omitted but all other staining steps kept unchanged. Right, image showing undetectable fluorescence when the primary antibody was omitted but all other staining steps were left unimpaired. (**B**) Similar representations as shown in A except that the secondary antibody was conjugated to Abberior Star Red (Ab.St.RED). Scale bar, 500 nm. (**C**) Stacked column graph showing negligible fluorescence (∼0.1%) when either the SV (orange) or the primary antibody (magenta) was omitted during the staining (n= 5 independent experiments for both Ab.St.RED and AF594). (**D**) Representative images (fire) of Synaptotagmin 1(Syt1) and specified VTs in purified chromaffin granules (CGs). Syt1, which is endogenously present on CGs was used as a marker for CGs. Scale bar=500 nm. (**E**) Stack column graph shows quantification of the number of puncta for the indicated combination of antibodies.

**Figure S4.**
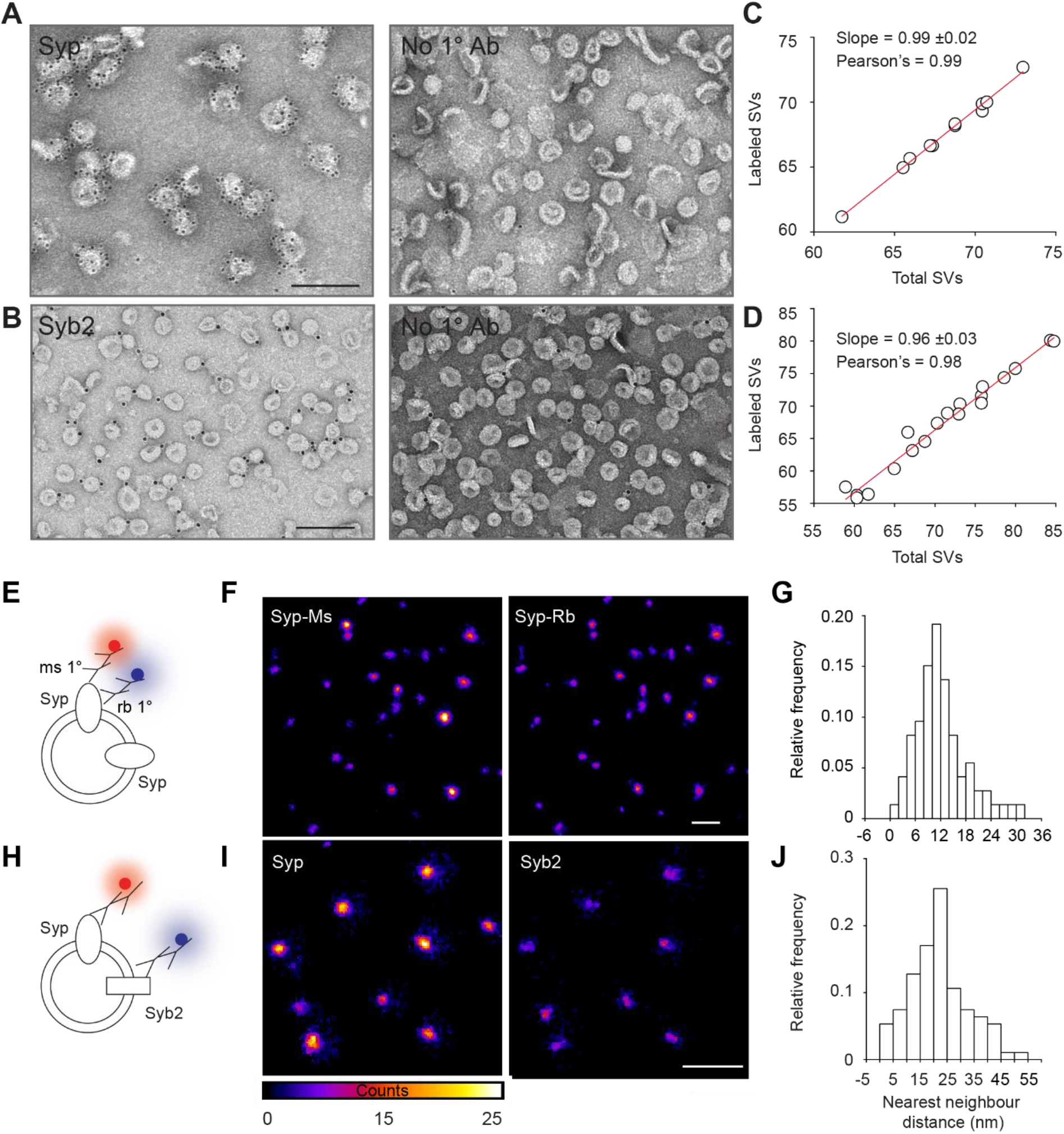
Immunogold electron microscopy (IEM) and two-color DyMIN STED reveals Syp and Syb2 as markers for total number SVs. (**A**) Representative immunogold electron microscopy (IEM) image of purified SVs labeled against Syp. (**B**) Representative IEM image of purified SVs labeled against Syb2. The images on the right represent the negative controls that lack the respective primary antibodies. Anti-Syp antibody typically labeled all small vesicular profiles with ∼10 gold particles per vesicle. Anti-Syb2 antibody labeled ∼96 % of vesicles but mostly with only 1 gold particle per vesicle. Scale bars in A and B, 100 nm. (**C**) and (**D**) Scatter plots quantifying the labeled SVs for Syp and Syb2 in IEM. Total SVs (X-axis) are plotted against gold-labeled SVs (Y-axis) in individual images. Data points were fitted to a linear model (n = 2 experiments). Labeling of almost all SVs by Syp and Syb2 antibodies in IEM, which exhibits limited labeling efficiency, increases the likelihood that the two antibodies stain all SVs in DyMIN STED imaging, which exhibits supra-optimal conditions because of in-solution immunolabeling. (**E**) Illustration depicting two-color labeling of the same protein, Syp, in purified SVs using mouse and rabbit anti-Syp primary antibodies. (**F**) Representative DyMIN STED images (fire) of the two fluorescence channels. Scale bar, 500 nm (**G**) Histogram showing the distribution of nearest neighbor distances (NND) of Syp-Ms puncta in Syb2-Rb channel. (**H**) Illustration depicting double immunolabeling of SVs against Syp and Syb2. (**I**) Representative DyMIN STED images showing single vesicles expressing Syp (left) and Syb2 (right). Scale bars in F and I, 500 nm (**J**) Histogram showing the distribution of nearest neighbor distances of Syp puncta in Syb2 channel. Note the distances between centers of all puncta in G and H is <55 nm, which allowed setting up of a threshold for the distance between the coordinates of two fluorescence puncta for the determination of colocalization of proteins on the same SVs (n = 4 experiments, >14,000 SVs in each channel).

**Figure S5.**
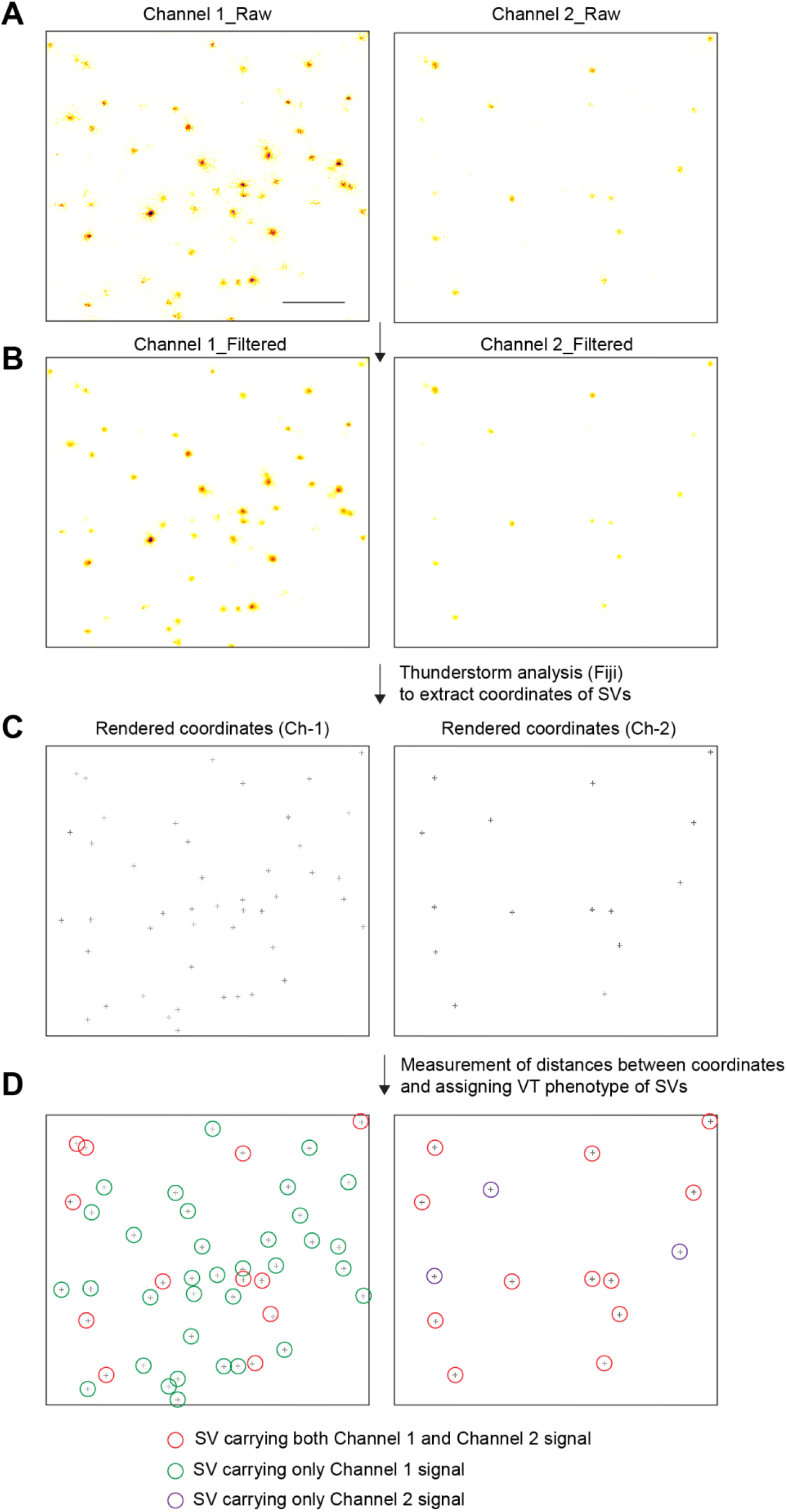
Overview of single vesicle colocalization analysis workflow. (**A**) Representative raw images (inverse LUT) of an example two-channel DyMIN STED acquisition of single vesicles labeled against two SV proteins (Channel 1 and Channel 2). Scale bar, 500 nm. (**B**) Same paired images as above are shown following median filtering with a radius of 1 pixel (15 nm). (**C**) Rendering of the above paired images showing extracted coordinates of detected puncta using Thunderstorm analysis, an open source Plugin in Fiji. (**D**) The distance between the coordinates of any two puncta and in paired images was calculated. Two proteins were considered as colocalized on the same vesicle if the distance between the coordinates is less than 50 nm (see Supplementary Materials for details), yielding VT phenotype of detected SVs and their corresponding percentage composition.

**Figure S6.**
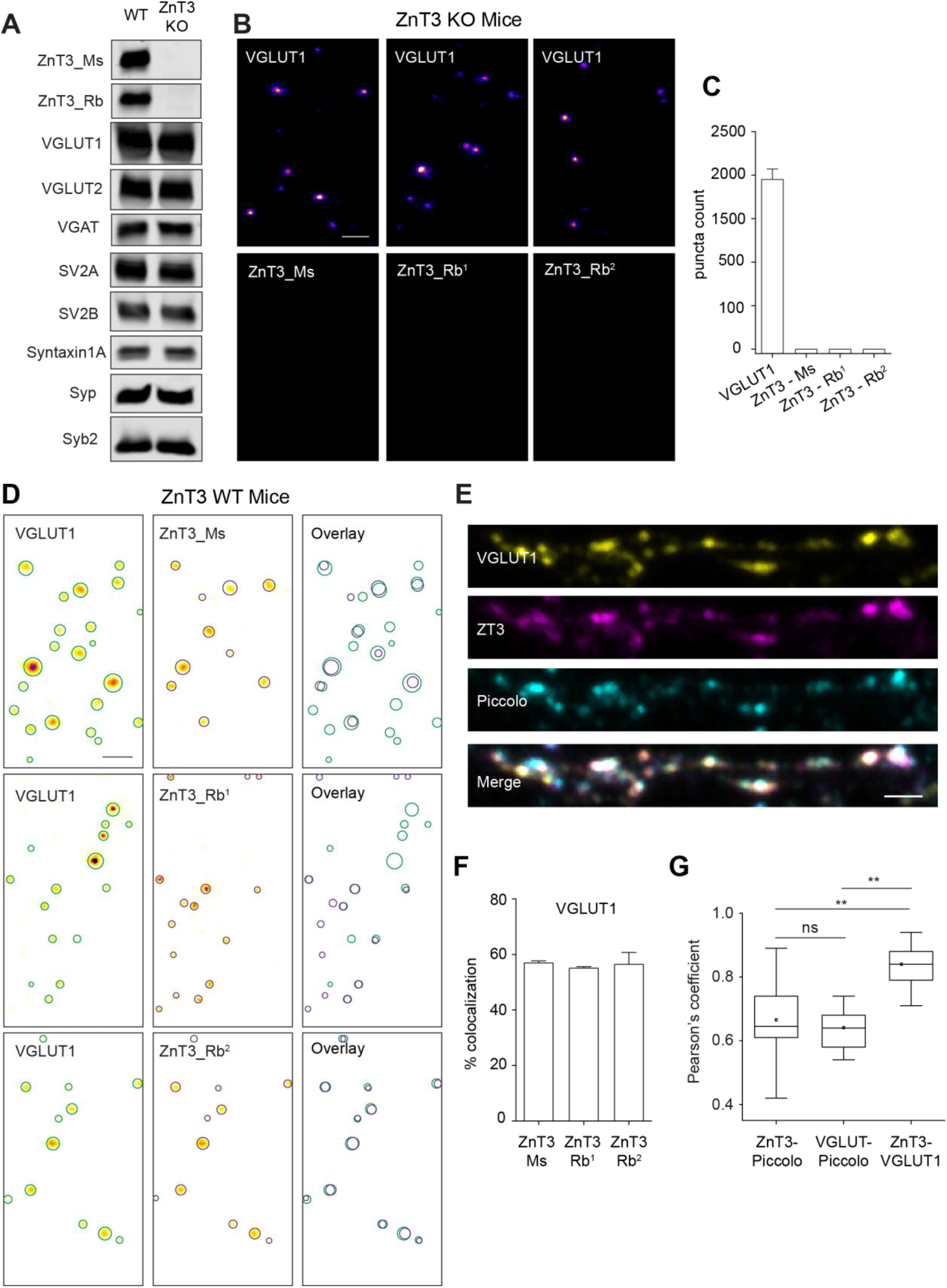
Validation of robust VGLUT1 and ZnT3 colocalization in mouse preparations and hippocampal culture neurons. (**A**) Western blot detection of ZnT3 and other selected SV proteins in wild type (WT) and ZnT3 knock out (KO) SV preparations (LP2). Both mouse and rabbit antibodies against ZnT3 (Ms- SySy-197011; Rb^1^- SySy- 197003) were not able to detect any ZnT3 in the KO derived sample, thus validating the specificity of ZnT3 antibodies used in the current study. However, there was no apparent difference in the expression of other proteins tested in the KO mice, including VGLUT1. (**B**) Representative images of single VGLUT1 and ZnT3 SVs purified from ZnT3 KO mice for the specified antibodies (Ms- SySy-197011; Rb^1^- SySy- 197003; Rb^2^- Alomone Labs-AZT-013). (**C**) Bar graph showing quantification of VGLUT1 and ZnT3 SVs in the KO (**D**) Representative two-color DyMIN STED images (inverse LUT) of VGLUT1 and ZnT3 (specified antibodies) channels in the wildtype. Circles portray SV area derived by a 2D Gaussian fit on VGLUT1 (green) and ZnT3 (magenta) puncta. The overlay shows SV circle profiles of the both VGLUT1 and ZnT3 channels. Scale bar, 200 nm. (**E**) Representative confocal images of dendritic segments of hippocampal culture neurons labeled against VGLUT1, ZnT3 and Piccolo, a presynaptic active zone marker. Merge shows colocalization of the three proteins. Scale bar, 1 µm. (**F**) Bar graph quantifying degree of colocalization between VGLUT1 and ZnT3 SVs in the WT mice. (**G**) Box plot showing significant spatial correlation between VGLUT1 and ZnT3 signals in comparison to their colocalization against Piccolo (**P<0.01 n=3 experiments, unpaired t test). No significant difference observed for VGLUT1 and ZnT3 colocalization with respect to Piccolo (p=0.72, ns, unpaired t-test). Data in C and F are mean ± SEM.

**Figure S7.**
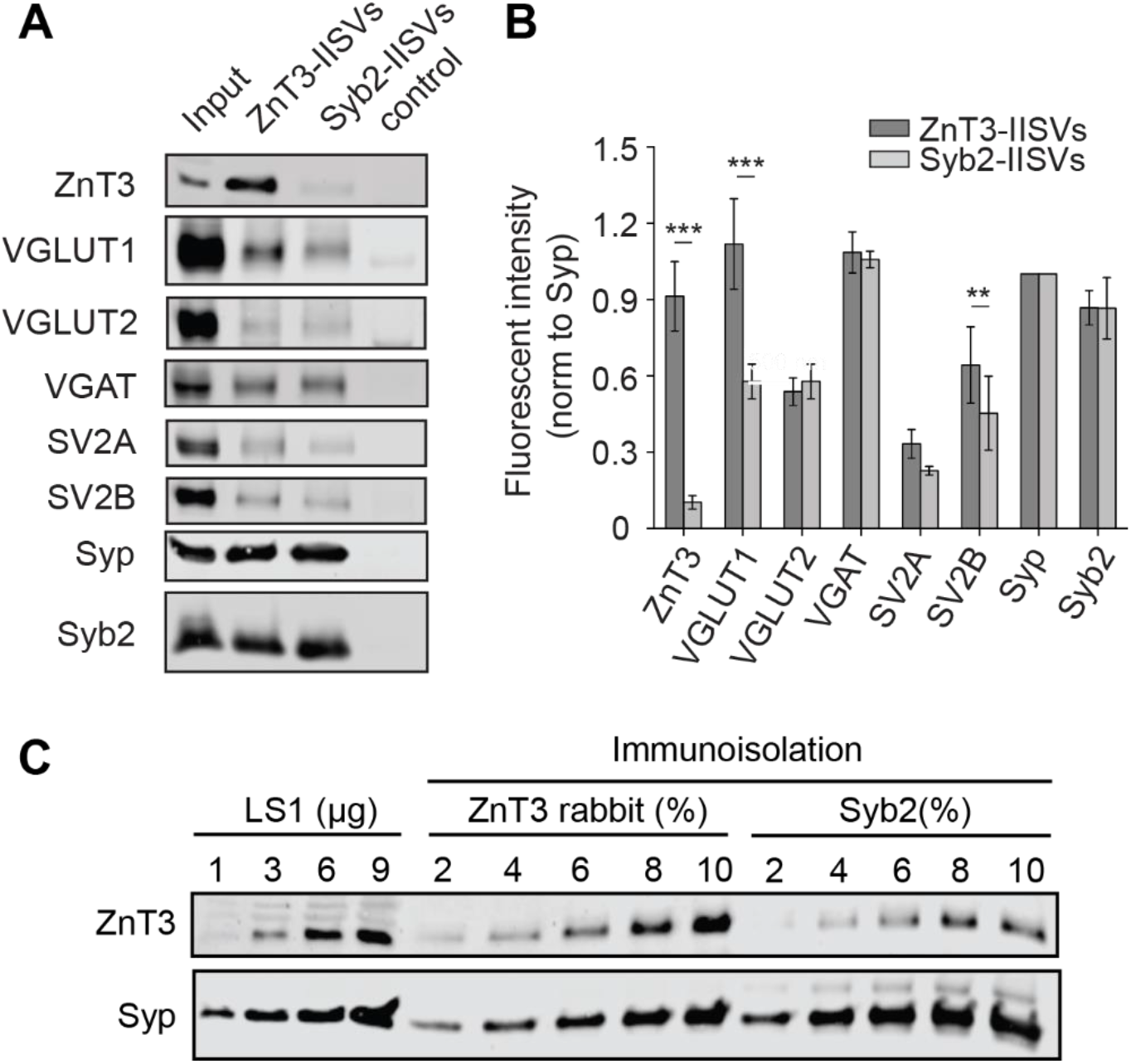
Immunoisolated ZnT3 SVs show enrichment for VGLUT1 and SV2B. (**A**) Immunoblot showing differential enrichment of selected proteins in immunoisolated SVs (IISVs) using specific antibodies against ZnT3 and Syb2 (substitute for all SVs). Note that only VGLUT1 and SV2B are enriched in ZnT3 immunoisolated vesicles. (**B**) Summary bar graph showing significant enrichment of VGLUT1 and SV2B in ZnT3 immunoisolated vesicles (\*\*\**P*<0.001, \*\**P<*0.05, n = 3 experiments, unpaired *t* test) (**C**) Western blot analysis showing determination of loading volume for enrichment assay shown in B and C using Syp as reference. The experiment was performed on LS1, ZnT3 and Syb2 immunoisolated samples for different loading volumes and detected against ZnT3 and Syp. The loading volume corresponding to equivalent Syp bands between ZnT3 and Syb2 samples were taken for western blot enrichment assay. Data are mean ± SEM.

